# Longitudinal multi-omic evaluation of biomarkers of health and ageing over smoking cessation intervention

**DOI:** 10.1101/2025.08.29.673135

**Authors:** Chiara Maria Stella Herzog, Charlotte D. Vavourakis, Bente Theeuwes, Elisa Redl, Christina Watschinger, Gabriel Knoll, Magdalena Hagen, Andreas Haider, Hans-Peter Platzer, Umesh Kumar, Sophia Zollner-Kiechl, Michael Knoflach, Nora Gibitz-Eisath, Stefan Öhler, Verena Lindner, Anna Wimmer, Tobias Greitemeyer, Peter Widschwendter, Sonja Sturm, Hermann Stuppner, Birgit Weinberger, Alexander Moschen, Alexander Höller, Wolfgang Schobersberger, Christian Haring, Martin Widschwendter

**Affiliations:** European Translational Oncology Prevention and Screening (EUTOPS) Institute, Universität Innsbruck, Milser Str. 10, A-6060 Hall in Tirol, Austria; Institute for Biomedical Aging Research, Universität Innsbruck, A-6020 Innsbruck, Austria; Department of Internal Medicine 2, Faculty of Medicine, Johannes Kepler University Linz, 4020 Linz, Austria; Institute for Sports Medicine, Alpine Medicine and Health Tourism (ISAG), University Hospital/Tirol Kliniken, A-6020 Innsbruck, Austria; Institute of Pharmacy/Pharmacognosy, Universität Innsbruck, A-6020 Innsbruck, Austria; Department of Neurology, Medical University Innsbruck, Innrain 52, A-6020 Innsbruck, Austria; VASCage - Centre on Clinical Stroke Research, Adamgasse 23, A-6020 Innsbruck, Austria; Department of Neurology, Hochzirl Hospital, Hochzirl 1, Zirl, Austria; Labordiagnostic St. Gallen West AG, St. Gallen, Switzerland; Suchthilfe Tirol, Innsbruckerstraße 85, A-6060 Hall in Tirol, Austria; Department of Gynaecology and Obstetrics, Landeskrankenhaus Hall, Tirol Kliniken, A-6060 Hall in Tirol, Austria; Institute for Psychology, Universität Innsbruck, A-6020 Innsbruck, Austria; Department of Gynaecology and Obstetrics, University Hospital Ulm, Prittwitzstrasse 43, 89081 Ulm, Germany; Danube Private University, Steiner Landstraße 124, 3500 Krems an der Donau, Austria; Division for Nutrition and Dietetics, University Hospital Innsbruck, A-6020 Innsbruck, Austria; Institute of Public Health, Medical Decision Making and Health Technology Assessment, Department of Public Health, Health Services Research and Health Technology Assessment, UMIT TIROL – University for Health Sciences and Technology, A-6060 Hall in Tirol, Austria; Digital Health Information Systems, Center for Health & Bioresources, AIT Austrian Institute of Technology, Graz, Austria; UMIT Tirol - Private University for Health Sciences and Health Technology, ISAG, A-6060 Hall, Austria; Tirol Kliniken, Anichstrasse 35, A-6020 Innsbruck; Department of Women’s Cancer, UCL EGA Institute for Women’s Health, University College London, Medical School Building, Room 340, 74 Huntley Street, WC1E 6AU, London, UK; Department of Women’s and Children’s Health, Karolinska Institutet, Stockholm, Sweden

## Abstract

Smoking is one of the single most important preventable risk factors for cancer and other adverse health outcomes ^1,2^. Smoking cessation represents a key public health intervention with the potential to reduce its negative health outcomes ^2–4^. While epidemiological, cross-sectional, and individual longitudinal ‘omic’ or biomarker studies have evaluated the impact of smoking cessation, no study to date has systematically profiled molecular and clinical changes in several organ systems or tissues longitudinally over the course of smoking cessation that could allow for more detailed assessment of response biomarkers and the identification of interindividual differences in the recovery of physiological functions. Here, we report the first human longitudinal multi-omic study of smoking cessation, evaluating 2,501 unique single or composite features from 1,094 longitudinal samples. Our comprehensive analysis, leveraging over half a million longitudinal data points, revealed a profound effect of smoking cessation on epigenetic biomarkers and microbiome features across multiple organ systems within 6 months of smoking cessation, alongside shifts in the immune and blood oxygenation system. Moreover, our multi-omic analysis provided unprecedented granularity that allows for identification of new cross-ome associations for mechanistic discovery. We anticipate that data and an interactive app from the Tyrol Lifestyle Atlas (eutops.github.io/lifestyle-atlas), comprising the current study and a parallel study arm evaluating the impact of diet on biomarkers of health and disease, will provide the basis for future discovery, biomarker benchmarking in their responsiveness to health-promoting interventions, and study of individualised response group, representing a major advance for personalised health monitoring using biomarkers.

## Main text

Smoking is a major risk factor for many chronic diseases and represents the leading modifiable risk factor for cancers worldwide ^1^. Efforts to aid smoking cessation or reduction of tobacco smoking are tantamount for public health as these interventions have the potential to alleviate the burden of smoking-related adverse health outcomes such as cancer ^2–4^. Studying molecular changes over the course of smoking cessations holds the potential to better understand the normalisation of homeostatic processes, reduction of disease risk, and temporal dynamics of smoking- and smoking-related disease biomarkers. While separate studies have evaluated changes in epigenetics ^5,6^, metabolome ^7^, microbiome ^8–10^, and other ‘omics’ (e.g., mutation load ^11^) individually using cross-sectional or longitudinal studies of current and former smokers, no study to date has comprehensively profiled multiple omics across several systems over the course of a smoking cessation intervention in humans. Here, we report on the first systemic multi-omic intervention study of smoking cessation in healthy women, assessing 2,501 unique single or composite features in 1,094 longitudinal samples, resulting in over half a million longitudinal data points. We collected detailed epidemiological information and longitudinally profiled clinical and functional features, immune populations, blood, cervical, and buccal methylation, faecal and saliva microbiome, and urine and saliva metabolome, allowing for analysis of both within omes and multi-omic integration. We observe a gradual but significant normalisation of epigenetic, immune, and microbiome features over 6 months of smoking cessation, and identify new associations between epigenetic and other systemic features, potentially increasing their future interpretability. Our results provide the first compendium of smoking cessation-associated molecular changes and data from the Tyrol Lifestyle Atlas ^12^ may be used as a new resource to benchmark the responsivity of biomarkers associated with smoking-related disease (e.g., epigenetic biomarkers). Together with findings from a parallel study on intermittent fasting, we anticipate that the Tyrol Lifestyle Atlas ^12,13^ will provide a basis for future mechanistic discovery of new biomarkers to monitor the individual effectiveness of smoking cessation and other health interventions.

### Study overview

Between April 2021 and February 2022, 42 female participants aged 30-60 were recruited into the TirolGESUND study smoking cessation arm. Baseline participant characteristics are shown in **Extended Data Table 1**. Detailed study information including participant flow charts are provided in a data and study description paper ^12^. A total of 24/42 (57.1%) participants completed the four follow up visits of the study, of whom 14 (58.3%) reported quitting smoking by month 6. Individuals who quit smoking exhibited a lower number of baseline cigarettes per day and smoking pack years. Individuals who dropped out of the study exhibited similar ages and BMI as individuals who did not drop out, but tended to have higher baseline fasting glucose values and a lower VO_2_peak (**Extended Data Table 1**). Biological samples for detailed molecular and cellular profiling were obtained every two months for six months while functional sports exams alongside detailed clinical blood measurements and vascular sonography were conducted at baseline and at six months (**Figure 1a**). Personal coaches recorded the number of weekly cigarettes and potential use of nicotine replacement tools (including gum, patches, or e-cigarettes), providing fine-grained information as well as allowing grouping of participants in higher and lower compliance, as defined by smoking cessation at the end of the study or not (**Figure 1b**). One participant each reported the use of nicotine gum and plaster, respectively. Two participants reported the use of heated tobacco products (‘heets’) while only one participant reported the use of e-cigarettes. The remaining participants did not report the use of any nicotine replacement products. Urine cotinine measurements were performed to confirm short-term nicotine abstinence.

**Figure 1.**
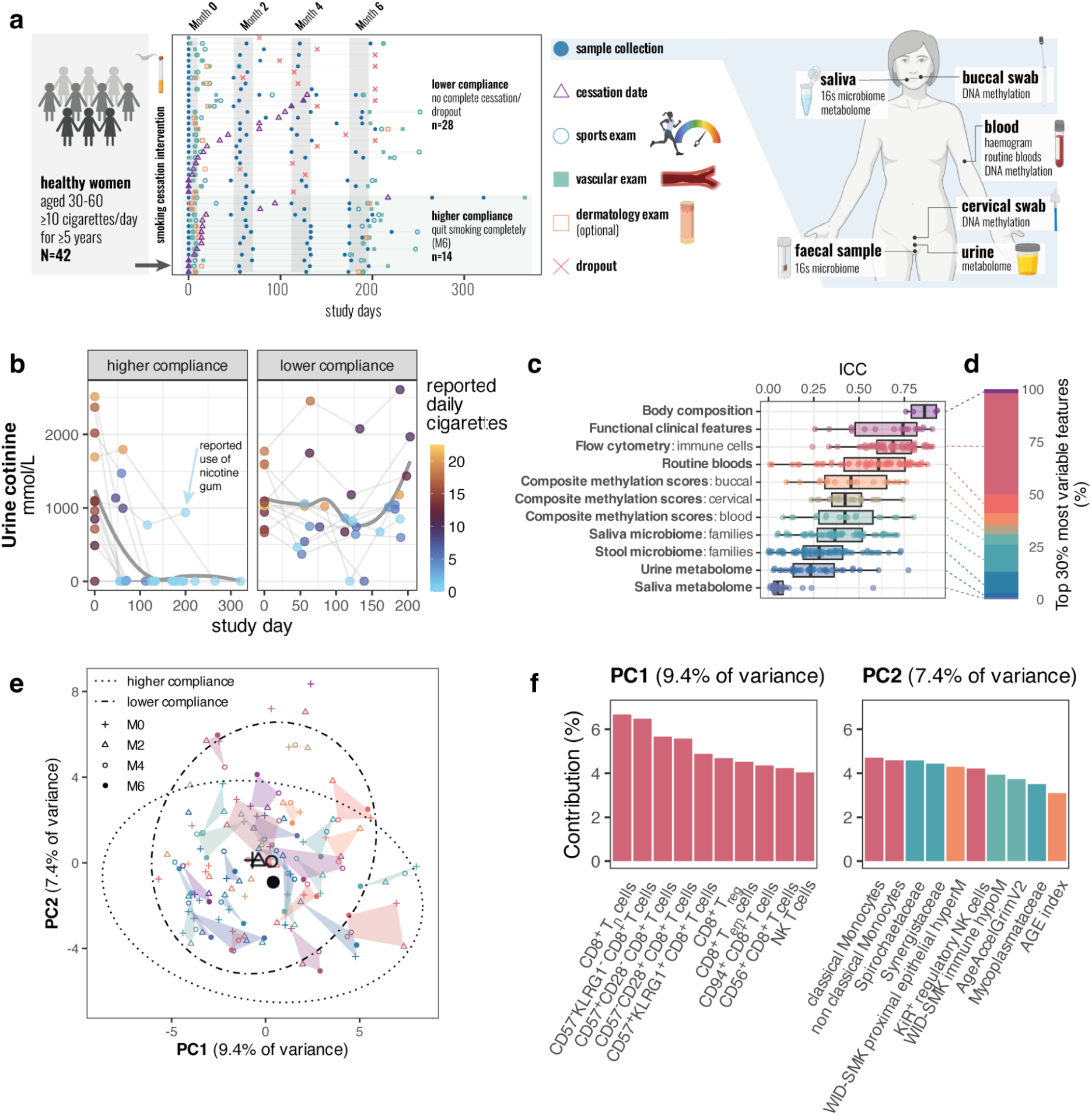
Study overview and clinical changes. **a** Diagram of study overview and collected samples. **b** Urine cotinine and self-reported cigarettes over the study days show agreement and allow for grouping of participants into higher and lower compliance groups. Regression line was fitted using loess method in ggplot. **c** Intra-class correlation (ICC) from linear mixed-effects models of normalised data, defined as proportion of the random variance out of the total variance. **d** Assays distribution of top 30% of most variable longitudinal data, including only features with more than 2 repeated measurements. Functional clinical measures were excluded as they were assessed twice instead of four times. **e** Principal components of the most personal 30% of features from Panel **d** (highest 30% based on ICC in features with 4 longitudinal measurements) over time. Shapes indicate visit (M0-M6), hulls indicate individuals, and outline ellipses show overall features in higher and lower compliant individuals (same grouping as in b). **f** Top 10 contributing features to principal components 1 and 2, coloured by assay (as in c-e). **Abbreviations**: ICC, intra-class correlation; M, month; PC, principal component.

The collection of longitudinal data allowed us to assess changes over time within and between individuals across several molecular and cellular features. We evaluated the variance explained by the participant structure using the intraclass correlation coefficient (ICC) from linear mixed-effects (LME) models to obtain an initial insight into the variation of different data types. Consistent with previous longitudinal human multi-omic studies ^14,15^, clinical laboratory tests and immune-related features were more personally distinct than other features (e.g., metabolomic measurement; **Figure 1c**). Decomposing the variance of the top 30% of personally variable features with 4 longitudinal measurements (assay sources shown in **Figure 1d**), we observed that the first two principal components (PCs) were driven by both inter-individual and intervention-associated features (time and compliance). PC1 was associated primarily with age and smoking pack years and its top contributors were immune cell populations (**Extended Data Figure 1a, b, Figure 1f**), while PC2 was changing over time (**Figure 1e**) and associated with previously described indicators of smoking (e.g., WID-SMK proximal epithelial hypermethylation, **Figure 1f**).

To study the longitudinal effects of smoking cessation in more detail, we next went on to characterise changes across clinical, epigenetic, immune cell profiles, metabolomic, and microbiome features performing both an *intention to treat* analysis and leveraging compliance groups (**Figure 1b**) to identify *per protocol* effects of smoking cessation (comparing to the partial reduction of daily cigarettes that also occurred in the lower compliance group). Detailed results of Wilcoxon tests and LME models adjusting for covariates are reported in **Extended Data Tables 2-7** and are available for interactive exploration on the Tyrol Lifestyle Atlas Data Portal (<u>eutops.github.io/lifestyle-atlas</u>).

### Clinical changes over the course of smoking cessation

Principal component (PC) analysis of the longitudinal clinical data of 24 individuals who completed the study revealed a small shift in principal components 1 and 2 (22.6 and 10.6% of variability, respectively) driven by contributors such as body mass index, body weight, and exercise capacity measures (PC1) and factors related to blood haematocrit, platelet volume, and haemoglobin (PC2) (**Figure 2a-c**), in line with previous reports of post-smoking cessation weight gain ^16^ and an association of smoking with increased haematocrit, platelets, and haemoglobin ^17,18^. Linear mixed-effects (LME) models confirmed these changes, including an increase in body mass index (BMI) and weight and an overall reduction in erythrocytes and haematocrit (**Figure 2d**), and demonstrated that higher compliance significantly exacerbated the increase in BMI and weight over time relative to the lower compliance group (**Figure 2d**), although overall changes in BMI were moderate (Δ in BMI of 0.36 kg/m^2^ (SD: 1.345) between M0 and M6, **Extended Data Table 2**). Participants exhibited an increase in their peak heart rate during ergometry (exercise exam) and a reduction in leukocytes, diastolic blood pressure, liver and kidney blood parameters (creatinine kinase and gamma-glutamyl transpeptidase), and surprisingly, in lung function parameters, compared to baseline (**Figure 2d**), although none of these were significantly modulated by higher compliance. Relative to the lower compliance groups, individuals who were highly compliant with smoking cessation exhibited reduced non-HDL and LDL at month 6 (**Figure 2d**). No significant alterations were observed in other clinical features, including vascular parameters (e.g., pulse wave velocity or intima-media thickness) or functional exercise capacity measures. Overall, participants who completed the study exhibited some clinical alterations, primarily associated with the blood oxygenation and clotting system, including erythrocytes, haematocrit, and platelets. We observed a small but significant increase in weight and BMI but did not detect any other major clinical benefits within 6 months, including lung function or exercise capacity, within the 6-month study window.

**Figure 2.**
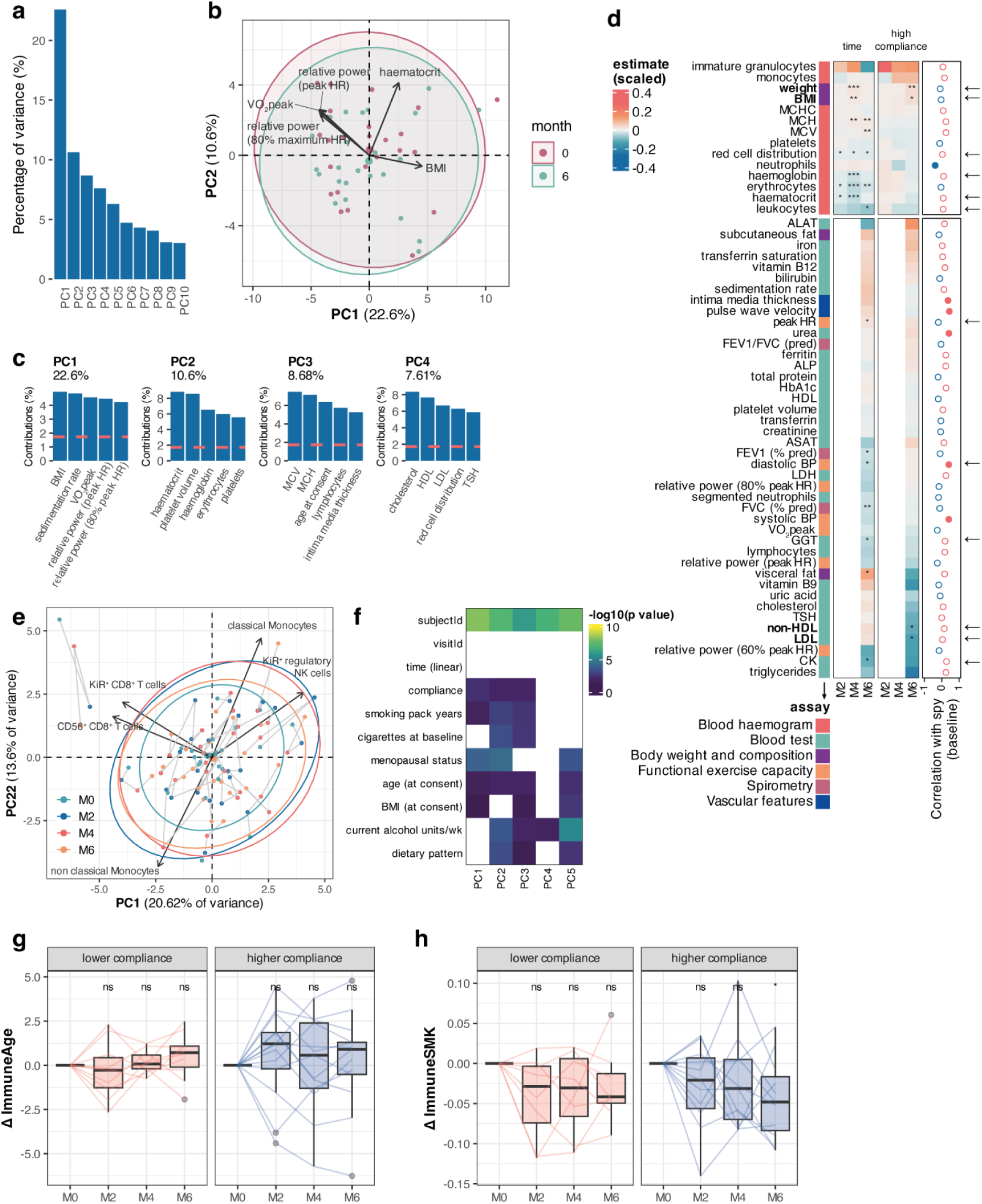
Clinical and immune changes over the course of the smoking cessation intervention. **a** Scree plot explaining percentage of variance of the top 10 principal components derived on clinical data from M0 and M6. **b** Principal components 1 and 2. **c** Top 5 contributors to the first four principal components. **d** Estimates from linear mixed-effects (LME) models of time or the interaction of time and high compliance (high compliance). Points on the right show correlation of the variable with smoking pack-years (spy) at baseline, and arrows indicate a significant modulation of the interaction of time and high compliance in the opposite direction. **d** Results from linear mixed effects (LME) models, estimating the impact of time overall (value ∼ age at consent + smoking pack years at consent + visitId + (1|subjectId)) and high compliance via an interaction model (value ∼ age at consent + smoking pack years at consent + visitId*compliance + (1|subjectId)). *, **, *** indicate p < 0.05, < 0.01, and < 0.001, from linear mixed-effects models, extracted via lmerTest, respectively. Correlation with age at baseline smoking pack year (spy) values is indicated on the right hand side (positive or negative in red and blue, respectively). Significant correlations at p<0.05 are indicated with filled circles. Changes that are going in the opposite direction of the correlation with smoking pack years at baseline are indicated with an arrow. **e** Biplot of the first two principal components of flow cytometry data on complete cases. **f** PCA heatmap of associations of the first five principal components with participant characteristics, computed either via Pearson correlation test for numeric values or Kruskall-Wallis test for factorial values. **g** Changes in ImmuneAge from baseline by compliance group. **h** Changes in ImmuneSMK by compliance group. p values in g and h are computed using paired two-tailed Wilcoxon tests, with * denoting p<0.05. Individual data points are shown wherever possible. Box plots correspond to standard Tukey representation, with boxes indicating median and interquartile range, and lines indicating smallest and largest values within 1.5 times of the 25th and 75th percentile, respectively. Individual data points are overlaid. No corrections for multiple testing were carried out. **Abbreviations**: PC, principal component; CRP, c-reactive protein; BMI, body mass index; MCHC, mean corpuscular haemoglobin concentration; MCH, mean corpuscular haemoglobin; MCV, mean corpuscular volume; ALAT, alanine aminotransferase; HR, heart rate; FEV1, forced expiratory volume (1s); FVC, full vital capacity; pred, predicted; ALP, alkaline phosphatase; HbA1c, haemoglobin A1c ; HDL, high density lipoprotein; ASAT, aspartate aminotransferase; LDH, lactate dehydrogenase; BP, blood pressure; VO_2_peak, peak rate of oxygen consumption; GGT, gamma glutamyl transferase; TSH, thyroid stimulating hormone; LDL, low density lipoprotein; CK, creatinine kinase.

### Smoking cessation alters the immune system and boosts numbers of cytotoxic NK cells

Our exploratory analysis indicated that immune features were associated with age and smoking pack years (**Figure 1e, Extended Data Figure 1a, b**), in line with previous reports ^19–21^. Evaluating 50 immune cell populations in peripheral blood in more detail, we found that several populations were associated with smoking pack years at baseline at p<0.05, including (classical) monocytes, plasma cells, central memory cytotoxic T cells (CD8^+^ T_cm_) (positive) and naive cytotoxic T cells (CD8^+^ T_n_), cytotoxic NK cells, and CD57^-^ KLRG1^-^ CD8^+^ T cells (negative) (**Extended Data Figure 2a, b**), although none remained significant after adjustment for multiple testing. Principal component analysis on individuals who completed the study (n=24) indicated a shift in overall immune cell features over time although none of the first five principal components were significantly associated with time point of visit (visitId) (**Figure 2e, f**), and suggested that monocytes and NK cells may be among top contributors to changes in immune cell proportions over time (**Figure 2e**). LME models confirmed a significant increase in several NK cell-related populations over time, including cytotoxic and regulatory NK cells and NK T cells, albeit no significant modulation by compliance was observed (**Extended Data Figure 2c, d**). In addition to NK-related immune population, several other cell populations changed. Highly compliant individuals also exhibited a significant reduction in classical monocytes, and CD94^+^ regulatory NK cells over time relative to individuals who were less compliant (**Extended Data Figure 2c, d**). To interpret changes in immune cell composition in the context of ageing and smoking cessation, we evaluated an immune population-based estimate of age (ImmuneAge) ^13^ and an immune population-based predictor of current smoking (ImmuneSMK; see **Supplementary Figure 1 and Methods**). We did not observe any significant changes over time in the ImmuneAge measure, suggesting that smoking cessation does not modulate age-related changes in immune populations (**Figure 2g**), but found a significant reduction of ImmuneSMK in highly compliant individuals at month 6 (**Figure 2h**). Taken together, our data demonstrated that although immune populations are personally distinct features with a high ICC (**Figure 1c**), smoking cessation significantly impacts several immune populations, including the adaptive immune system system, significantly reducing an immune-based estimator of recent smoking within 6 months of smoking cessation (**Figure 2h**).

### Significant reduction of some epigenetic features associated with smoking and biological ageing after smoking cessation

The epigenome, in particular, DNA methylation (DNAme), of buccal, cervical, and blood cells is known to be profoundly impacted by non-heritable factors such as smoking ^22,23^ and ageing ^24^ and may integrate the impact of heritable and non-heritable factors ^25^, representing an ideal biomarker to estimate the effect and potential long-term consequences of health interventions. Several biomarkers have been developed based on DNAme analysis in these relatively non-invasive surrogate samples, to indicate the current rate of ageing (DunedinPACE ^26^), presence ^27,28^ and/or future disease risk ^29,30^ of distant cancers (WID indices) and other diseases such as diabetes ^31^, healthspan (PhenoAge ^32^), mortality (GrimAge ^33^), or causal age-related changes (CausAge ^34^), amongst others. However, less is known about the stability of DNA methylation biomarkers and their responsivity to health-promoting interventions such as smoking cessation that may represent an important criterion for their translational application ^35^. Here, we therefore characterised the dynamics of key epigenetic biomarkers of smoking, disease risk, and ageing, leveraging longitudinal Infinium MethylationEPIC DNA methylation data from three sample types (buccal, blood, and cervical samples) to perform a comprehensive benchmarking of their association with smoking and sensitivity to change.

Our exploratory analysis indicated that an epigenetic biomarker of smoking was among the top personal features and contributed to a change in PC2 over time (**Figure 1e**). Variation in the overall top 5% variable CpGs on the Infinium MethylationEPIC array after quality control was primarily driven by sample composition and influenced by other covariates, including age, smoking pack years, compliance, and others (**Figure 3a, Extended Data Figure 3a, b**). With the exception of a small but significant increase in estimated immune cell fraction (based on a methylation-based deconvolution algorithm, see Methods) in buccal samples at month 4, no significant changes in sample composition were observed over time (**Figure 3b**). We thereafter focused on relevant composite smoking, ageing, or cancer biomarkers in each sample type (**Figure 3c-f**).

**Figure 3.**
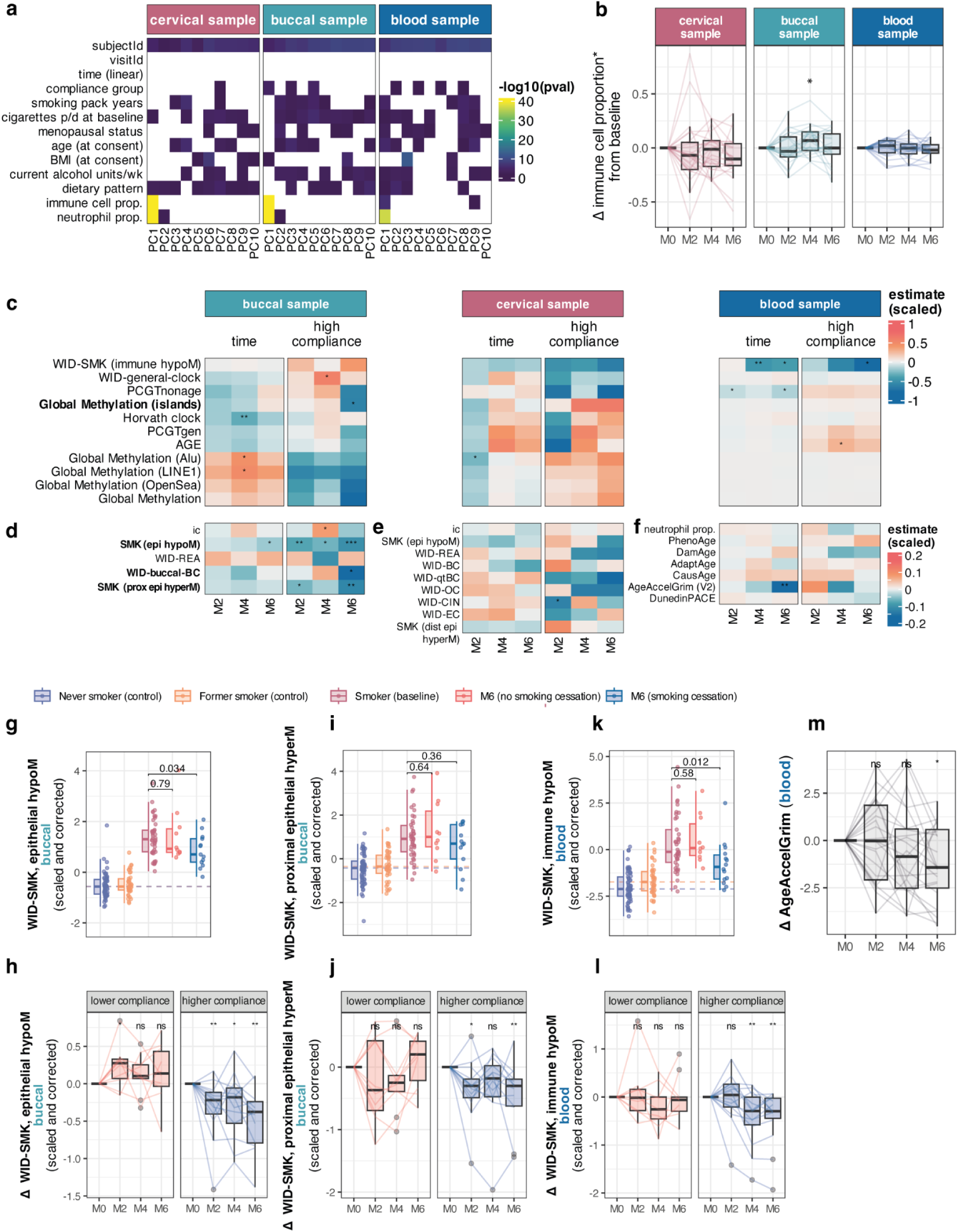
Partial recovery of smoking- and ageing-associated epigenetic features after smoking cessation. **a** Heatmap of features significantly associated with the first 10 principal components of the top 5% variable CpGs on the IlluminaHumanMethylation EPIC array. **b** Changes in immune cells (cervical or buccal) or neutrophil proportion (blood) in longitudinal samples of all individuals (intention to treat). **c** Heatmap of linear mixed-effects model estimates for time or interaction of time with high compliance in indices shared across multiple tissues, **d** buccal-specific indices, **e** cervical-specific indices, or **f** blood-specific indices. Share in c-e is the same (−1 to 1), whereas **f** has a different scale (−0.2 to 0.2). **g** WID-SMK epithelial hypoM in buccal samples of never, former, current smokers (all at baseline) or at 6 months, split by compliance. **h** Longitudinal changes in WID-SMK epithelial hypoM, grouped by compliance. **i** WID-SMK proximal epithelial hyperM in buccal samples of never, former, current smokers (all at baseline) or at 6 months, split by compliance. **j** Longitudinal changes in WID-SMK proximal epithelial hyperM, grouped by compliance. **k** WID-SMK immune hypoM in blood samples of never, former, current smokers (all at baseline) or at 6 months, split by compliance. **l** Longitudinal changes in WID-SMK immune hypoM, grouped by compliance. **m** Longitudinal changes in GrimAge Acceleration in blood samples. *, p<0.05, **, p<0.01, in LME model (c-f), unpaired two-sided Wilcoxon test test (g-k) or paired two-sided Wilcoxon test compared to baseline (m-l). Box plots correspond to standard Tukey representation, with boxes indicating median and interquartile ranges and whiskers indicating ± 1.5 times interquartile ranges. Individual data points are shown where possible.

We initially validated recently described cell-specific biomarkers of smoking. Previously developed scores were transformed to always exhibit the same directionality in smokers versus non-smokers, i.e., an elevation, and were centred around the mean using baseline data z-scaling. Scores in baseline samples of current smokers (this study), former smokers with several quit-years or never smokers (parallel study ^13^) confirmed that recently-described WID-SMK biomarkers ^22^ behaved as anticipated (**Extended Data Figure 3c**) and additionally revealed a preferential association of some cell-specific biomarkers with recent or acute measures of smoking dose (urine cotinine and buccal WID-SMK epithelial hypoM) while others exhibited a preferential association with cumulative measures of exposure (estimated smoking pack years and buccal WID-SMK proximal epithelial hyperM) (**Extended Data Figure 3d**). Assessing smoking-related biomarkers in buccal, blood and cervical samples over time revealed that some, but not all, biomarkers were significantly changed over time: WID-SMK epithelial hypoM and WID-SMK proximal epithelial hyperM (buccal) and WID-SMK immune hypoM (blood) were reduced at month 6 overall, with a significantly greater reduction in individuals with higher compliance compared to individuals who did not quit smoking (**Figure 3c-l**). Notably, in contrast to immune hypoM in blood and epithelial hypoM in buccal samples, buccal WID-SMK proximal epithelial hyperM was significantly reduced in longitudinal paired tests but not significantly different to baseline values of overall smokers, indicating larger individual variability and potential slower recovery (**Figure 3i, j**), as perhaps expected due to its association with cumulative but limited association with more recent measures of smoking exposure (**Extended Data Figure 3d**). Biomarkers of smoking in cervical samples were not significantly altered after 6 months of smoking cessation, although epithelial hypoM, immune hypoM and distal epithelial hyperM all exhibited negative estimates in LME models (**Figure 3c, e**). The WID-CIN index ^27^, a biomarker for cervical cancer and pre-neoplasias, was significantly reduced in the high compliance group relative to the lower compliance group at month 2, and several other biomarkers were modulated by smoking (**Figure 3e**). Lastly, we also observed a reduction in GrimAge acceleration, an established biomarker of biological age and mortality risk ^33^ (**Figure 3m**), although this was not modulated by higher compliance (**Figure 3f**). Investigating individual components of GrimAge we found a significant reduction of DNAmPACKYRS, in line with an impact of smoking cessation on smoking-related composite scores (**Extended Data Figure 3e**).

We lastly explored the impact of nicotine replacement on smoking-related biomarkers in blood and buccal samples. Two participants who quit smoking switched to e-cigarettes or nicotine gum, respectively exhibited a strong reduction in blood immune hypoM (**Extended Data Figure 3f**), while only the e-cigarette but not the gum user exhibited a reduction in the buccal epithelial hypoM score, a signature possibly associated with nicotine detoxification and metabolism ^22^. This finding is in line with the observation that the nicotine gum user also exhibited consistently high levels of cotinine in urine (**Figure 1b**). Conversely, the gum user exhibited the strongest reduction of the WID-SMK proximal epithelial hyperM score, a signature based on methylation in many genes associated with carcinogenesis and possibly predictive of lung cancer risk ^22^, while the e-cigarette user exhibited a smaller reduction (**Extended Data Figure 3f**).

Our comprehensive benchmarking of several relevant epigenetic biomarkers of smoking, disease risk, and ageing reveal a normalisation of smoking-associated epigenetic changes and a reduction of GrimAge acceleration, driven primarily by an effect on the DNAmPACKYRS module, while they show no differences in several other biomarkers of ageing, cervical sample smoking biomarkers, and disease (**Figure 3c–f**), indicating a diverse responsivity of biomarkers across tissues to smoking cessation. Notably, longitudinal data in an individual who switched to e-cigarette use resulted in a strong reduction in immune smoking-related changes in blood and epithelial tobacco detoxification signature. Given the limited sample size in the current study to explore the impact of e-cigarette use in more detail, future studies with a larger sample size are warranted to explore the longitudinal effects of e-cigarette use as smoking cessation aids on biomarkers of smoking and cancer.

### Metabolome of saliva and urine exhibits high variability yet reveals distinct smoking cessation-associated changes

The low intra-class correlation coefficients (ICCs) of saliva and urine metabolome data sets (**Figure 1c**) suggested they may be less stable within an individual compared with other features and rather be driven by short-term changes prior to sampling, such as dietary intake, although sample collection at home was conducted using a standardised protocol (first-morning urine; saliva prior to brushing teeth). The urine metabolome exhibited a slightly higher ICC than the salivary metabolome. Principal component analyses aligned with the ICC and revealed that the first five principal components of the saliva metabolome, accounting for to 52.9% of variance, were not associated with subjectId, while some of the first five principal components of the urine metabolome, accounting for 46.6% of the variance, were (**Extended Data Figure 4a-d**). Surprisingly, within the classes of confidently identified metabolites, no individual metabolite was significantly associated with either smoking pack years (data not shown) or current number of cigarettes in either baseline saliva and urine metabolites (with the exception of Cotinine that was not considered in the current analysis; **Figure 1b**; **Extended Data Figure 4e-h**), possibly stemming from a large variability in metabolites in combination with a relatively low number of individuals.

Evaluating longitudinal changes in the saliva metabolome of individuals who completed the study, we nonetheless observed significant reductions of thymine, taurine, lysine, hydroxyacetone, and acetoin, comparing baseline and follow-up visits (**Figure 4a-c**). The change from baseline in trimethylamine N-oxide was significantly less pronounced in individuals with higher compliance, and individuals with higher compliance surprisingly exhibited higher saliva ethanol values than lower compliant individuals (**Figure 4a, e, f**), possibly stemming from differences in dietary habits. The higher presence of isopropanol was more unusual as it is typically not found in the human body and may be linked to exposures to certain environmental sources or substances. In urine, a significant increase in tartrate and glucose, and 2-hydroxyisobutyric acid, alongside reductions in taurine, lactic acid, trimethylamine, choline, 3-methylhistidine, acetic acid, betaine, and phosphocreatinine were observed over time (**Figure 4g-k**). High compliance significantly modulated levels of 2-hydroxyisobutyric acid, lactic acid, indoxyl sulfate, 4-hydroxyphenyl acetate, and glycine (**Figure 4g, h)**. Our analysis indicated that saliva and urine metabolomics overall exhibited highly variable features over time, but did show some distinct changes associated with the smoking cessation intervention, including biologically relevant metabolites previously associated with smoking. For instance, smoking or inhalation of heated air have been shown to increase blood lactate levels ^36^ while decreasing blood glucose levels ^37^, whereas cessation of smoking may explain reduced levels of lactate and increase levels of glucose in (blood and subsequently) urine, as observed in the current study. The reduction in salivary hydroxyacetone and acetoin was also noteworthy as these metabolite are present in e-cigarette liquids and has been found to be associated with the cytotoxic ^38^ or respiratory adverse effects ^39^ of e-cigarette aerosols, yet none of our participants reported smoking e-cigarettes at the start of the study. The increase in tartrate (M4 only) was interesting as it has been suggested to be a biomarker of moderate wine consumption ^40^. Lastly, choline and trimethylamine lie within the same metabolic pathway via anaerobic microbial conversion ^41^, and both metabolites exhibited a significant reduction in urine, suggesting significant changes in this pathway over the course of the intervention.

**Figure 4.**
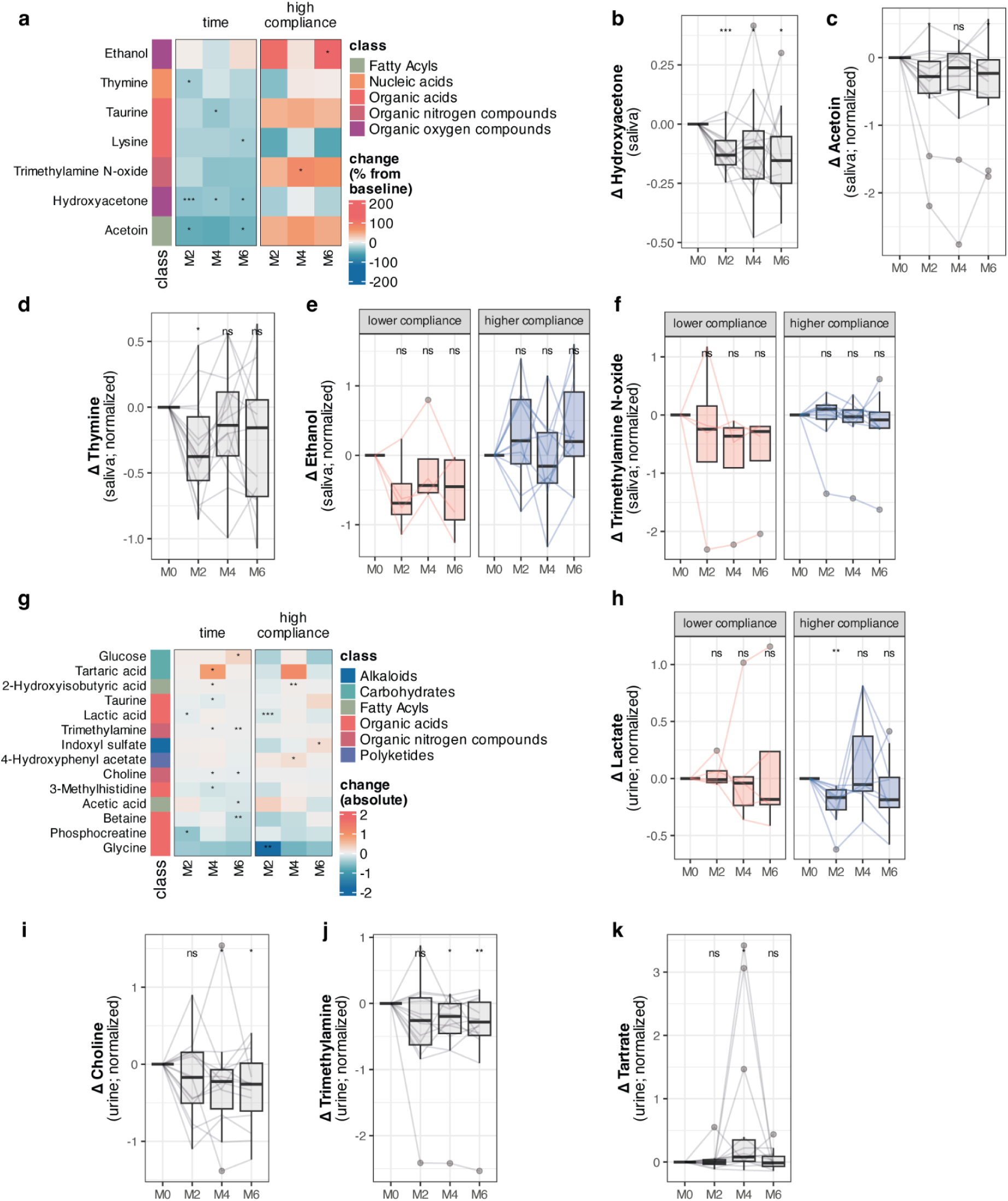
Metabolome alterations over the course of smoking cessation. **a** Heatmap of significant salivary metabolome changes from baseline (as %) and corresponding Wilcoxon test p values in complete cases overall (‘time’ column, paired two-sided Wilcoxon test) or comparing the differences from baseline in high versus lower compliance groups (‘high compliance’ column, unpaired two-sided Wilcoxon test). The left annotation indicates the metabolite class according to RefMet. *, p<0.05; ***, p<0.001. **b** Paired longitudinal changes in salivary hydroxyacetone, **c** acetoin, **d** thymine, **e** ethanol (by compliance group) and **f** trimethylamine N-oxide (by compliance group). p values are derived from paired two-sided Wilcoxon tests compared to baseline values. **g** Heatmap of absolute changes in the urine metabolome and corresponding Wilcoxon test p values in complete cases overall (‘time’ column, paired two-sided Wilcoxon test) or comparing the differences from baseline in high versus lower compliance groups (‘high compliance’ column, unpaired two-sided Wilcoxon test). The left annotation indicates the metabolite class according to RefMet. *, p<0.05; **, p<0.01; ***, p<0.001. **h** Paired longitudinal changes in lactate (by compliance group), **i** choline, **j** trimethylamine, and **k** tartrate. p values in b-f and h-k are derived from paired two-sided Wilcoxon tests compared to individual baseline values. Box plots correspond to standard Tukey representation, with boxes indicating median and interquartile ranges and whiskers indicating ± 1.5 times interquartile ranges. Individual data points are shown where possible.

### Smoking-associated changes in the oral and gut microbiome are partially alleviated upon cessation

To identify microbiome features associated with smoking, we compared 16S rRNA gene amplicon sequence profiles of the saliva and stool samples from the smoking cessation study participants to matched profiles from never smokers (parallel study ^13^, n=24, BMI range=24.7-27.7; **Extended Data Figure 5a-b**). Saliva samples from smokers at baseline and the never smokers differed significantly in beta diversity estimated using the Aitchison distance (PERMANOVA, p-value < 0.05), demonstrating that smoking has a profound effect on the oral microbiome (**Figure 5a, Extended Data Table 8**). Smoking cessation gradually restored the diversity in the six months cessation period, with no difference in beta-diversity between the ex-smokers and never smoker group after six months, a significant difference between smokers and the never smoker group, and a near-significant difference between the smokers and the ex-smokers (pairwise PERMANOVA, p-values adjusted, alpha = 0.05). In contrast, beta diversity measured in stool samples was not significantly different between the smoking status groups at any given time point (**Figure 5b**).

**Figure 5.**
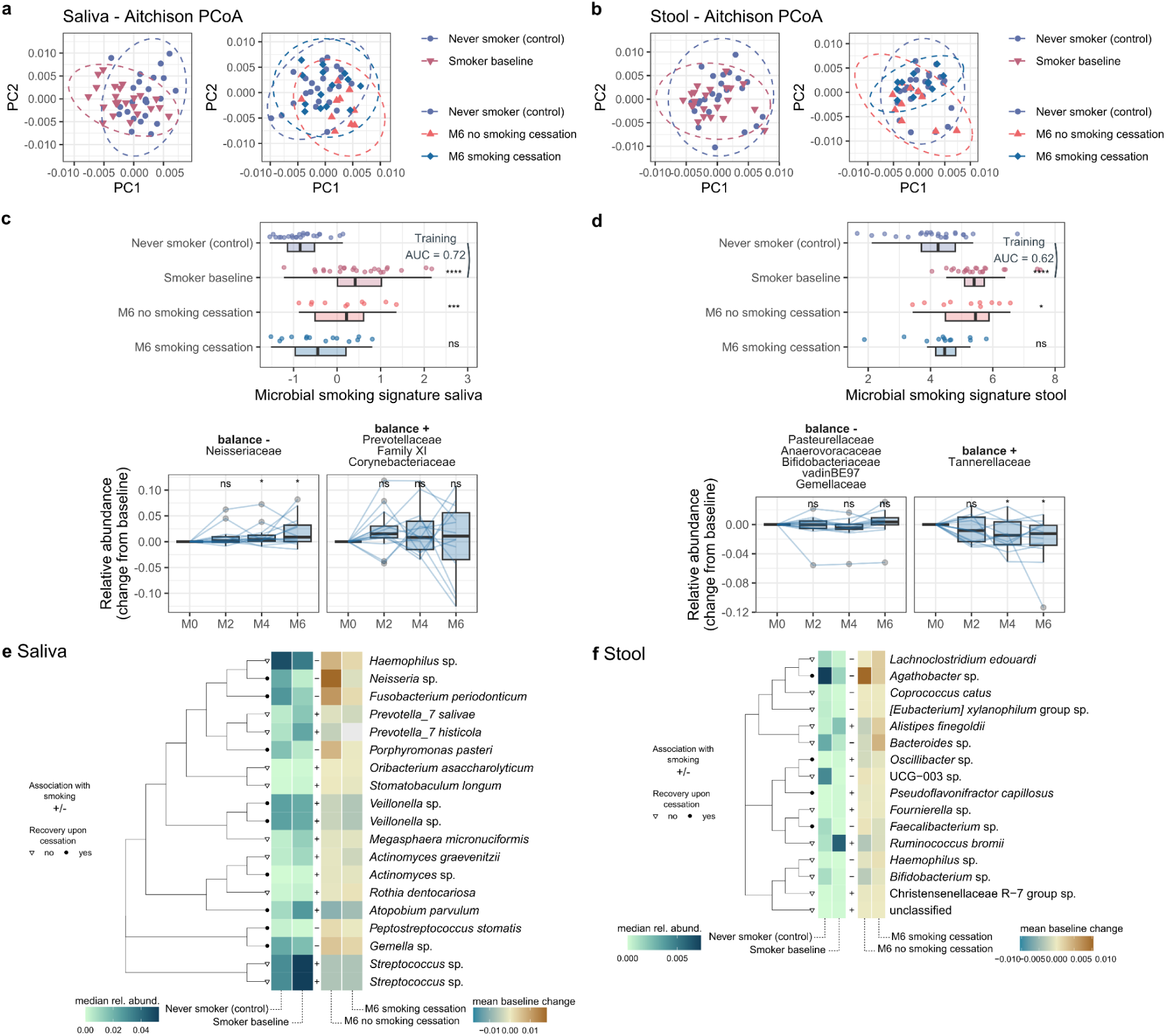
Smoking- and smoking cessation associated features in the oral and gut microbiome. **a-b** Principal coordinate analysis of Aitchison distances calculated for never-smoker, former-smoker and smoker groups. **c-d** Top panels: balanced, smoking-associated microbial signatures trained on family-level baseline data of never-smokers and smokers (AUC = the mean cross-validated area under the curve). Bottom panels: change in relative abundance of the microbial families contributing negatively or positively to each balance. Stars indicate the significance levels comparing medians of each group with that of the never-smokers (paired two-sided Wilcoxon signed-rank tests, with ns = non significant; * = p value<0.05; *** = p value<0.001). **e-f** Left side of the heatmap: median relative abundance of detected smoking-associated ASVs in never-smoker and smoker-groups and the (+/-) direction of the smoking-association. Right side of the heatmap: mean baseline change of the respective at month 6 in relative abundance in the former-smoker and smoker groups. Cladograms in the middle depict the phylogenetic relation between the ASVs. Tip annotation indicates whether a smoking-associated ASV recovers in abundance upon cessation (full circles; p value<0.05) or not (empty triangles). Box plots correspond to standard Tukey representation, with boxes indicating median and interquartile ranges and whiskers indicating ± 1.5 times interquartile ranges. Individual data points are shown where possible.

Next, we constructed balanced family-level microbial signatures associated with smoking for both body sites (**Figure 5c-d**). Training signatures on the never smoker and smoker data at baseline resulted in a mean cross-validation AUCs of 0.72 and 0.62 for the saliva and stool samples respectively, in line with our earlier observation that smoking has a more profound impact on the oral microbiome than on the stool microbiome. Smoking cessation led to partial recovery of both microbiomes, resulting in no significant difference in the signatures between never smokers and the ex-smokers after six months (p-values ≥ 0.05; **Figure 5c-d**, top panels). The relative abundance of *Neisseriaceae* in the oral microbiome was negatively associated with smoking and significantly increased over time in the higher compliance group (**Figure 5c**, bottom panel). In the gut microbiome, the relative abundance of *Tannerellaceae* was positively associated with smoking and significantly decreased over time with smoking cessation (**Figure 5d**, bottom panel). The abundances of the remaining families associated with smoking in both signatures were not modulated by smoking cessation. Longitudinal LME modelling confirmed the significance of the differential abundance of *Neisseriaceae* and *Tannerellaceae* with smoking cessation in the higher compliance group **(Extended Data Figure 5c-f)**. In addition, several other families were detected to change over time in the high compliance group. Of those, *Atopobiaceae* and *Comamonadaceae* were also significantly associated with smoking at baseline in the oral microbiome (Wilcoxon signed-rank test comparing never smokers, p-value<0.01; **Extended Data Figure 5e**) and respectively decreased and increased in abundance upon high compliance to the smoking cessation (LME higher compliance p-value < 0.001 for the time estimate at month 6; **Extended Data Figure 5c**).

Finally, to investigate the effect of smoking and smoking cessation at lower taxonomic levels, we first selected amplicon sequence variants (ASVs) with at least 90% and 50% prevalence in the saliva and stool samples, respectively (**Figure 5e-f**). From the 19 features that associated significantly with smoking in the oral microbiomes at baseline (Wilcoxon signed-rank test comparing never smokers, p<0.05), 8 were modulated after six months of smoking cessation (LME higher compliance p<0.05 for the time estimate at month 6; **Figure 5e**). Most notable was the decrease of two *Veillonella* spp. and the increase of *Porphyromonas pasteri* upon complete smoking cessation only. These species might be indicative of decreased biofilm formation ^42^ and improved periodontal and dental health ^43,44^. Interestingly, *Fusobacterium periodonticum* was found to be negatively associated with smoking and significantly increased in abundance upon cessation. Although this species was originally isolated from severe periodontitis lesions ^45^ and the more common *F. nucleatum* is an opportunistic periodontal pathogen associated with disease and oral cancer ^46^, the clinical relevance of *F. periodonticum* remains unknown. Amongst the remaining smoking associated ASVs whose abundance was not modulated by cessation, two high abundance *Streptococcus* spp., *Prevotella histicola*, *P. salivae*, *Megashpaera micronuciformis* and *Actinomyces graevenitzii* had the most distinct relative abundance between smokers and never smokers. In the stool only 4 out of 16 smoking associated ASVs showed signs of recovery after six months of smoking cessation (**Figure 5f**). The most remarkable changes observed upon cessation included an increase in the abundance of species within *Agathobacter* and a *Faecalibacterium*, two beneficial short-chain fatty acid producing genera ^47^. On the other hand, *Ruminicoccus bromii*, a specialist resistant starch degrader ^48^, was strongly associated with smoking, but did not significantly decrease in abundance after six months in the smoking cessation group. Notably, *Desulfovibrionaceae*, implicated in choline and trimethylamine metabolism ^41^, were not significantly altered by the intervention.

### Baseline predictors of smoking cessation success

We previously observed that baseline values of smoking pack years and cigarettes were slightly different between individuals that successfully quit smoking and those that did not or dropped out (higher compliance, lower compliance, and dropout in **Extended Data Table 1**), in line with previous observations that heavy smokers may exhibit lower success for smoking cessation interventions ^49^. We were therefore interested in systematically exploring the association of baseline values with smoking cessation success (yes, no, or dropout), and changes in two smoking-associated biomarkers at month 6 (WID-SMK epithelial hypoM, detoxification, and WID-SMK proximal epithelial hyperM, associated with potential cancer risk prediction). Smoking intervention success (grouped) was with current cigarettes at baseline (p=0.03), in line with observations in **Extended Data Table 1**. Some additional associations were observed for changes in methylation predictors. However, no feature remained significant after adjustment for multiple correction testing. Associations are shown in **Extended Data Table 9**.

### Multi-omic longitudinal integration uncovers systemic inflammatory changes

Our data, for the first time, enabled the integration of systemic multi-omic profiling over the course of 6 months of smoking cessation, enabling new discovery of cross-ome associations. **Figure 6a** shows an overview of timings of significant changes overall (intention to treat) in individuals that completed the study. For integrative analysis, we initially applied MEFISTO (Method for the Functional Integration of Spatial and Temporal Omics data ^50^) which revealed that, as expected, the first 11 factors were associated with subjectId and thus personally distinct, but some multi-omics factors were associated with participant and intervention details. Surprisingly, the first two factors were strongly correlated with each other (**Extended Data Figure 6a**). Factor 3 showed a significant association with compliance and values were significantly increasing over time, particularly in individuals with higher compliance to the smoking cessation intervention (**Extended Data Figure 6b, c**). Among the top 30 features contributing to factor 3 was the decrease in WID-SMK proximal epithelial hypermethylation, alongside several other epigenetic biomarkers of smoking, smoking-related cancer risk (WID-CIN, WID-buccal-BC), and ageing or mortality (Horvath clock, AgeAccelGrim V2) in blood, buccal, and cervical samples (**Extended Data Figure 6d**). We also found several saliva, but not gut microbiome families, contributing strongly to this factor, for example the decrease in Bifidobacteriaceae, Lactobacillaceae, Synergistaceae and Veillonellaceae. Interestingly, factor 3 was significantly increasing over time in individuals with higher compliance, but also showed a significant change at month 2 in lower compliance individuals (**Extended Data Figure 6c**). This finding is in line with the clinical observation that many individuals in the lower compliance group reported attempts to quit during the early phases of the study, with few self-reported cigarettes (**Figure 1b**), but eventually resumed smoking. These data also indicated a multi-omic response to smoking cessation, involving multi-tissue methylation changes and saliva microbiome alterations.

**Figure 6.**
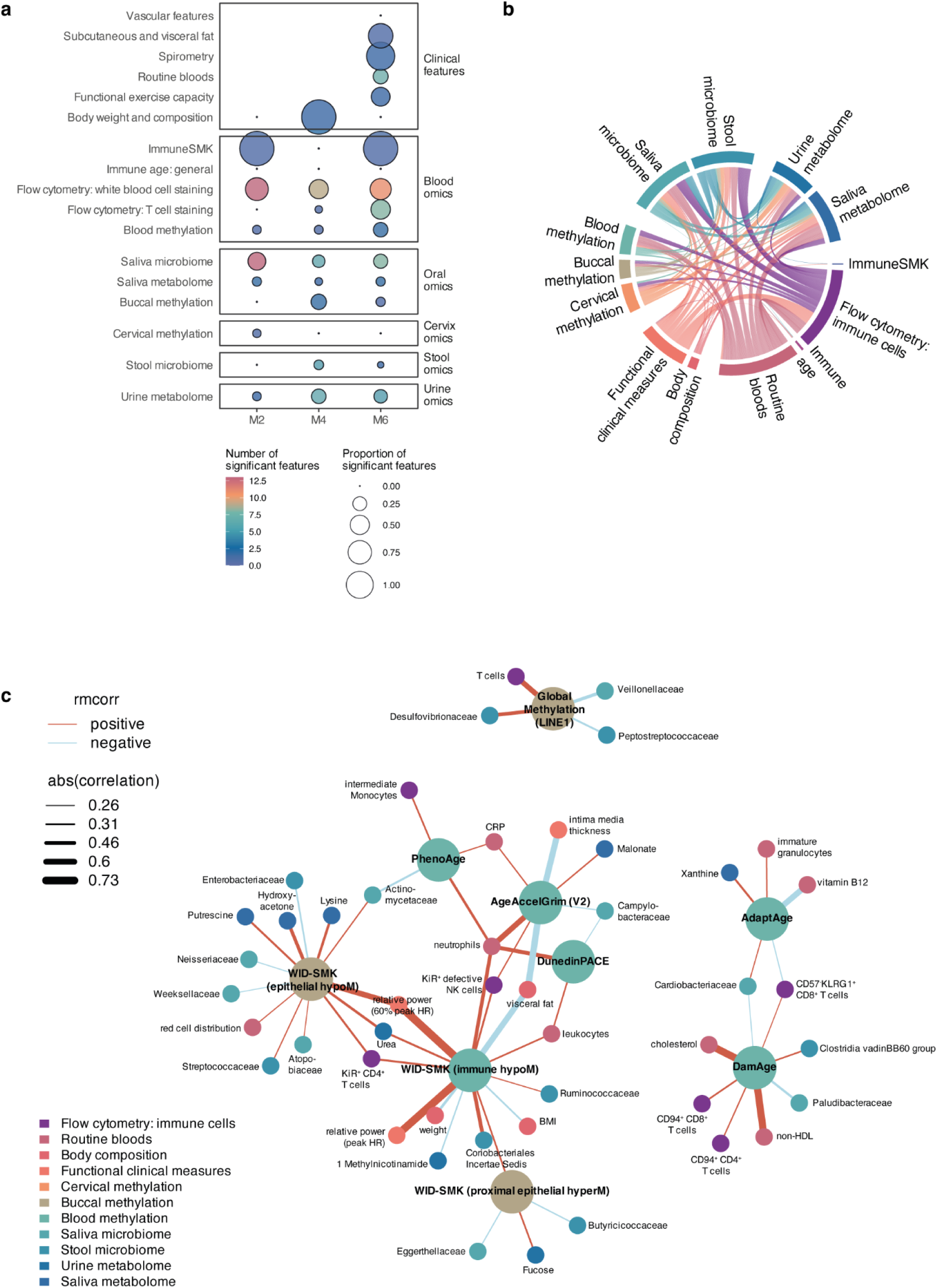
Integrative analysis of pathways across individuals. **a** Overview diagram visualising timing and number of changes across different omic and clinical features at p<0.05 in linear mixed-effects models (time estimate) or paired two-sided Wilcoxon tests (metabolome data only). **b** Chord diagram of significant cross-ome associations at p<0.01 for repeated measures correlation. **c** Network diagram of significantly associated features across key relevant biomarkers at p<0.01. Only immediate correlations are shown. Raw data are presented in Extended Data Table 9. **Abbreviations**: rmcorr, repeated measures correlation.

To further explore systemic associations of multi-omic features across the cohort, we leveraged the data to perform longitudinal repeated measures correlation analysis across 661 key features (clinical features, haemogram, flow cytometry, epigenetic biomarkers, metabolites, and microbiome at family level), focusing on cross-ome associations at a significance threshold of p<0.01 to favour discovery, which resulted in 753 significant pairwise associations of unique 321 features (**Figure 6b; Extended Data Table 10**). Repeated measures correlation analysis confirmed some previously known associations (e.g., c-reactive protein (CRP) and intermediate monocytes, **Extended Data Figure 7a**), but also facilitated discovery of new associations. For instance, we were interested in assessing associations of smoking- and ageing-associated epigenetic features. In **Figure 6c**, we show a correlation network constructed using features significantly correlated with several smoking- and ageing-related epigenetic features. Our analysis revealed that buccal WID-SMK proximal epithelial hypermethylation, located in genes associated with carcinogenesis and possibly associated with future lung cancer development ^22^, correlated positively with the WID-SMK blood immune hypomethylation score (r_rm_=0.23, p=0.0023), indicating that the two likely changed together, and urine fucose (r_rm_=0.28, p=0.017) but negatively with stool Butyricoccaceae (**Figure 6c**, **Extended Data Figure 7b-d**; r_rm_=-0.29, p=0.006). These findings are in line with prior literature: smokers exhibit increased risk of certain types of inflammatory bowel diseases, in particular Crohn’s disease ^51^, and there have been reports of a reduced abundance of Butyricoccaceae in individuals with inflammatory bowel conditions ^52–54^. At the species level, we observed a significant association of WID-SMK proximal epithelial hyperM with *Butyricicoccus*, a microbial species that has recently been trialled as a gut probiotic ^55^, as their produced metabolite butyrate is suggested to reduce intestinal inflammation ^56^. We note that overall, no significant change in Butyricicoccaceae was observed, so this observation warrants further investigation (**Figure 5**). Urine fucose has been suggested as a biomarker for liver disease and cancer ^57,58^. WID-SMK epithelial hypomethylation in buccal samples, a signature for the cellular response to the exposure to tobacco-related toxins ^22^, was associated with saliva hydroxyacetone, putrescine, and lysine, red blood cell distribution, and several saliva and stool microbiome families (**Figure 6c**). WID-SMK immune hypomethylation in blood samples was associated negatively with weight and BMI, and positively with relative power; these associations are likely to be specific to the current data set due to the observed increase in BMI. However, we also observed association with neutrophils, leukocytes, KiR^+^ T cells, several microbiome families, and urine 1-Methylnicotinamide (**Figure 6c**), that may likely point to biologically relevant associations in blood immune composition and other molecular changes.

We also explored associations of biomarkers of ageing with multi-omic features or biomarkers of smoking. PhenoAge, AgeAccelGrim and DunedinPACE were more closely associated with biomarkers of smoking, whereas AdaptAge and DamAge, causal biomarkers of ageing adaptations and damage, and buccal global LINE1 methylation, clustered further apart (**Figure 6c**). AdaptAge and DamAge exhibited opposing associations with Cardiobacteriaceae and CD57^-^KLRG1^+^CD8^+^ T cells, representing a subset of antigen-experienced T cells: DamAge was associated with an increased proportion of CD57^-^KLRG1^+^CD8^+^ T cells, whereas AdaptAge was associated with a reduced proportion, suggesting these cells may be involved in modulating the ageing process.

Taken together, these data provide a first insight into systemic associations of smoking and ageing-associated (epigenetic) biomarkers (**Extended Data Table 10**). Alongside other findings from a parallel study investigating the impact of intermittent fasting ^13^, these data are freely available in the Tyrol Lifestyle Atlas Data Portal (<u>eutops.github.io/lifestyle-atlas</u>) and will provide a basis for future mechanistic discovery and validation studies.

## Discussion

Here we describe the first longitudinal multi-omic intervention study of smoking cessation in healthy women over the course of smoking cessation, assessing over half a million data points from 42 individuals, 24 of whom completed the study.

Smoking cessation has been shown to alleviate the burden of tobacco-related disease ^2^. Cigarette smoking exhibits both acute and chronic or persistent effects ^59–61^ whose reversal likely occurs at different timescales. Acute effects as due to the presence of tobacco metabolites (cotinine) or elevated carbon monoxide levels may more rapidly normalise following cessation ^62^, whereas effects on (epi)genetic levels may take longer to reverse, if at all. The discrepancy between acute and long-term effects of smoking may explain why some cardiovascular and/or metabolic features ^63^ may more rapidly recover than cancer risk after smoking cessation ^4^. Epigenetic effects have been proposed to at least in part mediate some of the long-term health consequences of smoking ^23^. A recent study has proposed a ‘reversal’ of the genetic mutation load in bronchial tissue due to a gradual replenishment from a non-mutated pool of cells ^11^ whereas the situation for epigenetic alterations is less clear. DNA methylation features associated with smoking could on one hand be reversed by active de- or remethylation (no change in cell composition), or could be driven by a replacement of cells without such smoking-related alterations, in parallel to findings reported for DNA mutations in the bronchial epithelium, or a combination of the two mechanisms. Notably, tissues with faster turnover times may exhibit faster recovery. While the current study did not specifically investigate the mechanisms of DNA methylation normalisation following smoking cessation, our findings are in line with the replenishment hypothesis as we exhibit a significant reduction in smoking-associated signatures at 6 months in tissues with faster turnover (blood, buccal samples) but not in cervical samples, although future experiments, for instance single cell DNA methylation sequencing, will be required to investigate the mechanism in detail.

The data presented in the current study are largely in line with previous findings on metabolic and clinical changes. For instance, we observe a rapid change in the blood oxygenation system (**Figure 2d**) and several other biomarkers of cardiovascular fitness that may be related to acute changes in blood oxygenation, e.g., diastolic blood pressure, peak heart rate during exercise, and the clotting system (**Figure 2d**). However, we fail to observe a substantial improvement in clinical changes related to lung function, in contrast to some previous studies ^64,65^. We note that individuals were healthy at baseline and did not exhibit obvious compromises in lung function (**Extended Data Table 2**; FEV1 at 88.7%), though perhaps longer-term follow-ups may reveal a change in trajectories ^66^. Additional follow-ups are ongoing and may help to address this question in the future. In line with the literature, participants also experienced a small but significant increase in BMI despite dietary assistance over the intervention to mitigate the risk of weight gain. A recent study in mice identified dimethylglycine, produced by both gut microbiota and the host from dietary choline sulphate, as a key metabolite implicated in smoking cessation-induced weight gain^67^, warranting further investigation of a similar pathway in humans. Our study indicated significant smoking-cessation associated changes over time in urine concentrations of glycine, betaine and choline, which are intermediaries of this pathway. But since we did not observe a significant difference between stool samples from never-smokers and smokers at baseline in the predicted relative abundance of choline-o-sulfatase (EC 3.1.6.6), the key microbial enzyme of this pathway (results not shown), further concrete involvement of specific gut microbial enzymes remains to be investigated.

Recent work has demonstrated long-lasting effects of smoking in particular on the adaptive immune system and cytokine secretion ^61^. While we did not assess cytokine secretion in detail in the current study, we do observe significant and pronounced alterations in proportions of adaptive immune cells, in particular in highly compliant individuals, suggesting at least a partial restoration of smoking related alterations (**Figure 2h, Extended Data Figure 2c, d**). Future studies may explore data on cytokine production after LPS production in the current study, described in a separate data description paper ^12^, in more detail.

In addition to specific smoking-related epigenetic effects discussed above, epigenetic biomarkers also hold the potential to integrate exposures and monitor long-term disease risk, but their longitudinal responsivity to interventions remains largely unknown, hindering their translational potential ^35^. Here, we show that smoking cessation significantly reduces several epigenetic biomarkers (**Figure 3**), intriguingly those associated with smoking-related cancers (e.g., WID-CIN, a cervical cancer biomarker) and mortality (GrimAge). Our study also provides the first insight into systemic associations of (smoking-associated) biomarkers, identifying links to metabolites, immune cells, and microbes, that may aid their interpretability pending further mechanistic studies (**Figure 6b, c; Extended Data Table 10**). Our findings are available publicly on the Tyrol Lifestyle Atlas Data Portal, allowing for exploration and data visualisation.

Although data on additional covariates were available, in the current study we primarily focus on the effects of overall changes associated with smoking cessation. Our data also revealed significant associations of major principal components in many omic data types with other covariates, such as self-reported alcohol intake at baseline (**Figure 2, 3, Extended Data Figure 6**) that may warrant further investigation.

Our study has significant strengths, in particular the detailed clinical phenotypic and multi-omic profiling of several longitudinal biological samples. While previous studies had evaluated omic changes in never, former, and current smokers, including epigenetic features ^5^, faecal ^9,10^ and saliva ^8^ microbiomes, amongst others, and some longitudinal studies exist (e.g., DNA methylation in blood ^6^, metabolomics ^7^), to our knowledge no systemic multi-omic study describing changes during smoking cessation has been conducted to date. This study allowed us not only to assess changes within a multitude of relevant systems (saliva and faecal microbiome, functional clinical parameters, vascular system) and biomarkers (e.g., smoking and mortality using epigenetic biomarkers), but also, for the first time, associations between them. While the current study does not allow for confirmation of causality of these associations, we expect these findings will be highly informative for discovery and future mechanistic studies. Moreover, the current study validates recently described biomarkers of smoking ^22^ and identifies their links to acute or cumulative measures of smoking dosage (**Extended Data Figure 3d**). It would have been interesting to explore longitudinal changes in smoking-associated biomarkers when switching to e-cigarettes, but only one individual reported the use of e-cigarettes hence we could not feasibly assess this research question, although surprisingly, several reducing metabolites in saliva were associated with e-cigarette compounds (**Figure 4a, g**).

Our study also has limitations. Regrettably, we did not meet our initial target of 60 participants within the study recruitment window (April 2021-February 2022), which resulted in a smaller than anticipated sample size. Our study indicated that despite substantial mentoring and professional assistance with smoking cessation, including group sessions, the intervention resulted in only moderate success to achieve sustained smoking cessation (14/24 individuals, 58.3%). The drop-out rate of 42.9% (18/42) is in line with previously reported smoking cessation studies ^68^. Participants who dropped out reported higher numbers of daily cigarettes at baseline and tended to exhibit slightly worse fasting glucose levels and exercise capacity (VO_2_peak). Albeit the current study is small, the observation of a tendency for worse baseline health parameters may have important implications for future efforts of smoking cessation, as women with more unfavourable health determinants may benefit even more from mocking cessation. The current study only included women aged 30-60 with limited ethnic diversity, limiting its generalisability. For the purpose of the current study, we focused on discovery of new associations that will need to be explored in more detail in future studies, and thus the homogeneity of the study group was considered an advantage. While our study represents an important proof of concept, future studies should strive to include more diverse and larger study populations of diverse age groups, genders, and ethnicities to obtain a more complete picture of the molecular and systemic impacts of smoking cessation. A major limitation of our study is that we opted not to include a randomised no-intervention control group due to the significant participant effort required, and due to the fact that we did not deem it ethical to instruct participants to not quit smoking. Instead, we requested individuals who did not achieve smoking cessation to continue to provide samples, which allowed us to use this ‘lower compliance’ group as a nested comparison group (**Figure 1b**). While these findings need to be interpreted with caution, in particular given the small sample size, we believe the current study represents an important resource for multi-omic discovery due to its deep phenotypic and systemic multi-omic profiling. Future studies may also consider run in phases, which has been implemented in a more recent lifestyle intervention study for smoking cessation (<u>ISRCTN89257090</u>). Lastly, our analysis methods, such as linear mixed-effects models, rely on strong assumptions including relating data missingness, and we did not correct for multiple testing. We have mitigated the risk for false discoveries by leveraging longitudinal data wherever possible and exploring gradual changes over time. For instance, for many changes we observe show alterations in the same directionality over several visits, rendering us more confident in them (e.g., WID-SMK changes; **Figure 3**). We are committed to being fully transparent regarding our findings and provide our complete analysis code on Github (https://github.com/chiaraherzog/MultiOmics_SmokingCessation). Moreover, all results from intention to treat and per protocol analyses are provided in **Extended Data Tables** and on our Tyrol Lifestyle Atlas Portal (https://eutops.github.io/lifestyle-atlas/docs/app.html) for maximum reuse value, alongside sharing of anonymised raw data following an embargo period (details in ^12^).

Taken together, the current study presents human systemic molecular profiling at an unprecedented scale. We anticipate that this study can provide a platform for the discovery of new functional associations and hypothesis generation, as well as benchmarking of the longitudinal changes of biomarkers across a health-promoting intervention. For instance, future studies may investigate predictors of smoking cessation success or smoking-cessation associated weight gain in humans. While data presented in the current study are limited to specific omics and epigenetic biomarkers, the Tyrol Lifestyle Atlas (<u>eutops.github.io/lifestyle-atlas</u>) ^12^ additionally also features full epigenomic profiling of longitudinal samples, data on immune cell cytokines following stimulation, activity questionnaires, wearable data, and others that can be exploited for future studies.

## Methods

### Study overview

This study encompasses the TirolGESUND smoking cessation arm (<u>NCT05678426</u>). The TirolGESUND study has received ethical approval by the Ethics Committee of the Medical University of Innsbruck (Ethikkommission der Medizinischen Universität Innsbruck 1391/2020, 18.01.2021), Austria. The study is described in detail in a study description paper ^12^. In brief, healthy participants aged 30-60 who were current smokers, defined as smoking at least 10 cigarettes per day for the last five years, were recruited to participate in the study. All participants were healthy and free from current or former malignant, cardiometabolic, or psychiatric disease, and were not pregnant. Participants consented at baseline and completed several follow-up visits (month 2, 4, and 6, with optional month 12 and 18 follow-ups not included in the current analysis). Clinical measurements, samples, and questionnaire data were collected during visits or via digital tools at home.

### Clinical measurements

Functional clinical measurements were collected using spirometry and ergometry (exercise bike) at the Institute for Sports, Alpine Medicine and Health Tourism in Natters, Austria. Routine blood biochemistry was conducted using fasted samples while haemograms were performed fasted or unfasted. Vascular features and subcutaneous and visceral fat were measured using sonography and pulse wave velocity measurements at the Department for Neurology at the Medical University of Innsbruck, Austria.

### Methylation profiling

Methylation profiles were generated using the Illumina HumanMethylationEPIC version 1 in a high-throughput manner using liquid handling robots. Samples from the same individual were kept on the same beadchip, where possible, but positions of samples and visits were randomised to minimise batch effects. Data were preprocessed using a standardise pipeline (https://github.com/chiaraherzog/eutopsQC; previously reported e.g. in Herzog et al. ^22^) and thorough quality control was conducted to ensure no sample swaps occurred, including sample type analysis and SNP analysis of longitudinal samples. Some sample swaps could be identified and resolved based on SNP profiling and date of sample taking. Sample composition was estimated using methylation profiles using the EpiDISH algorithm (hierarchical EpiDISH for blood), using the centEpiFibIC reference matrix ^69^. Epigenetic biomarkers were computed using previously described coefficients or packages, including Biolearn (GrimAge ^70^).

### Immune cell profiling via flow cytometry

Peripheral blood samples were stored at room temperature for less than 24 h before isolation of peripheral blood mononuclear cells. Samples were stained and immune cell populations were profiled in detail by flow cytometry as previously described ^12^.

### Development of an ImmuneSMK classifier

We developed an immune population-based classifier of current cigarettes per day using baseline data (M0) from the smoking cessation and intermittent fasting study arms using all populations (normalised values) as input, exploring ridge, lasso, or elastic net penalisation in the glmnet R package (version 4.1.8). Elastic net penalisation exhibited the highest area under the curve to detect current smokers from non-smokers in baseline (training) data and was selected for the final model. Coefficients are shown in **Supplementary Figure 1a**. ImmuneSMK was then calculated using the supplied coefficients and longitudinally evaluated in follow-up samples.

### Saliva and urine metabolome profiling using nuclear magnetic resonance profiling and LC/MS

Saliva and urine metabolome data were generated using nuclear magnetic resonance profiling (see data paper ^12^). Spectra were annotated as previously described and manually checked. Raw values were normalised using Total Sum Scaling to account for differences in concentration. Cotinine quantitation was conducted via liquid chromatography-mass spectrometry (LC/MS) as described in our resource paper ^12^.

### Analysis 16S rRNA gene amplicons

DNA isolation, dual barcode sequencing ^71^ of the V4 variable region of the 16S rRNA gene and inference of amplicon sequence variants (ASVs) with DADA2 ^72^ was done as described in our resource paper ^12^ for all stool and saliva samples from the Smoking Cessation and Intermittent Fasting study arms collected within the TirolGESUND study. Automated taxonomic assignments by the SINA classifier ^73^ (version 1.7.2) against the SILVA database SSU Ref NR 99 (release 138.1) were further completed and curated down to the family-level. First, a maximum likelihood tree was constructed with FastTree ^74^ (-nt -gtr -gamma -spr 4 -boot 100) using a MAFFT ^75^ (--auto) alignment of all detected ASVs, which was then re-rooted between Archaea and Bacteria with the Newick utilities ^76^ and visualised using iTOL ^77^. Two ASVs that appeared as long branches in the tree and that were confirmed to originate from Eukaryotes (megablast against NCBI-nr) were removed. All remaining ASVs that could not be assigned down to the family-level were classified using Kraken2 ^78^ and the standard Kraken database (version 2.1.3). Full lineage classification for the resulting NCBI taxonIDs was extracted with TaxonKit ^79^. 106 out of 107 missing phyla labels successfully mapped to corresponding SILVA/SINA phyla (based on GTDB taxonomy), one ASV was manually assigned to the Bacteroidota (blastn, 91,6% sequence identity with *Prevotella pallens*). To complete the taxonomic table below the phylum-level, Kraken2 classifications were matched with corresponding SILVA/SINA classifications one level above using GTDB to NCBI taxonomy mappings [https://data.gtdb.ecogenomic.org/releases/latest/auxillary_files/, downloaded 17-10-2023]. Therefore, only the lowest level in manually curated assignment followed NCBI taxonomy. Separate ML trees were then constructed for ASVs belonging to Patescibacteria, Firmicutes (Bacillota), Bacteroidota, Actinobacteriota, Proteobacteria and any remaining phyla, using archaeal ASVs as an outgroup. Missing ASV taxonomic labels were then manually assigned, using an _unk suffix and the name of the highest confidently assigned taxon where needed. Finally, spurious ASVs that did not reach a relative abundance over 0.25 in any sample ^80^ were removed from the TirolGESUND saliva and stool microbiome datasets.

Only samples from the Smoking Cessation study arm and samples from 24 participants who had never smoked and who had the lowest body-mass index in the Intermittent Fasting study arm were used in downstream analyses. Differences in beta diversity between never-smoker, smoker and former-smoker groups were identified using the robust Aitchison distance ^81^ and Permutational Multivariate Analysis of Variance (PERMANOVA) ^82^ as implemented in the R package vegan ^83^ (version 2.6-4). Microbial signatures associated with smoking were constructed by training the selbal algorithm ^84^ (3-fold cross validation) on baseline count data summarised at the family-level for smokers and never-smokers. For prevalent ASVs with threshold 90% and 50% occurrence in samples collected at all time points for saliva and stool, respectively, smoking-association was inferred with Wilcoxon signed-rank tests (p value ≤ 0.05; R package stats version 4.2.0) using baseline relative abundance data from smokers and never-smokers. Relative abundance data of the ASV- and family-level microbiome features were used for longitudinal linear mixed modelling [as described above/below].

### Statistical analysis

We conducted both intention to treat and per protocol analysis, leveraging compliance categories (Figure 1) on complete cases. We performed both paired Wilcoxon tests as well as linear mixed-effects models that allowed us to additionally adjust for covariates (e.g., age or smoking pack years) and explore the interaction between time and compliance group. For standardised and harmonised data analysis, we leveraged the MultiAssayExperiment format ^85^ (MultiAssayExperiment R package version 1.28.0).

#### Intention to treat and per protocol analysis using Wilcoxon tests

Absolute clinical changes were evaluated using paired two-tailed Wilcoxon tests compared to baseline on complete cases for each feature (**Extended Data Table 2**), while differences in the change from baseline by compliance group were compared using unpaired two-tailed Wilcoxon tests (**Extended Data Table 4**). Longitudinal changes compared to baseline in the high compliance group only were computed using paired two-tailed Wilcoxon tests compared to baseline (**Extended Data Table 6**).

#### Intention to treat and per protocol analysis using linear mixed-effects models

In a second approach to assess the impact of time or the interaction between time and compliance on clinical and omic features, we fitted linear mixed effects (LME) models to fully leverage the longitudinal data, adjust for age at consent, and estimate the interaction between time since baseline and compliance. Briefly, each omic feature was normalised (thereafter referred to as ‘value’) and three models were run on the data, all containing fixed slopes and random intercepts. Model a) aimed to explore the impact of time overall on the value, accounting for age at consent, smoking pack years at consent, and visit (as factor). In the R lme4 package, the model was implemented as follows: value ∼ age at consent + smoking pack years at consent + visitId + (1|subjectId); Model b) evaluated the interaction between time and high compliance by extracting estimates for the interaction of visit (factor) and high compliance group (visitId*high compliance), accounting for age at consent, smoking pack years at consent. In the R lme4 package, the model was implemented as follows: value ∼ age at consent + smoking pack years at consent + visitId*compliance + (1|subjectId). Finally, model c) aimed to explore the impact of time and intervention group in highly compliant individuals by extracting the estimate for interaction of visit (factor) and interventionId (visitId*interventionId), accounting for age at consent and smoking pack years at consent. In the R lme4 package, the model was implemented as follows: value ∼ age at consent + smoking pack years + visitId + (1|subjectId[high compliance only]. We opted to code time as a categorical variable (visitId) to allow for non-linear effects (e.g., increase at month 2 with normalisation by month 6, as opposed to a gradual increase). Models with time as a continuous variable were also run (not shown, code available) and predominantly agreed with outputs from models with time as a categorical variable. Linear mixed models were run with the lme4 R package (version 1.1.35.1) and p values were extracted using the lmerTest R package (version 3.1.3).

#### Variance decomposition

To estimate between- and within-individual variation, we fitted linear mixed-effects models with fixed slopes and random intercepts, accounting for age at consent, smoking pack years at consent, and the interaction between visit (time as factor) and compliance. In R, the models were implemented as follows using the lme4 package (version 1.1.35.1): value ∼ visitId*compliance + age_at_consent + smoking pack years at consent + (1 | subjectId), where value represented the normalised feature. The intraclass correlation coefficient (ICC) was computed as the proportion of total variance explained by the subject structure, i.e. V_random_/V_total_, as extracted from linear mixed effects models using the get_variance function from the insight R package (version 0.19.8).

#### Association of baseline characteristics with smoking success or buccal methylation change

The association of baseline characteristics and smoking cessation success (higher compliance, lower compliance, or dropout), or change in WID-SMK proximal epithelial hyperM at month 6 compared to baseline was computed for each feature separately using Spearman correlation, Kruskall-Wallis, or Chi-Squared test, depending on data type (both continuous, continuous and categorical or both categorical, respectively). P values were adjusted using Holm-Bonferroni correction.

#### MEFISTO

We applied MEFISTO ^50^ to identify multi-omic longitudinal features in our data. Several models were trained: a) all data, regardless of intervention; b) all data or b) high compliance individuals only. Models were trained in the MOFA2 singularity environment in python (mofa2py v0.6.4). Model options were as follows: 10 factors; 200 iterations; n_grid=10; start_opt=50; opt_freq=50. The model trained on data from all individuals, regardless of compliance, exhibited the best model fit and was selected for further downstream exploration and association with covariates.

#### Repeated measures correlation analysis

We explore longitudinal associations of omic features using the rmcorr_mat function from the rmcorr R package (version 0.6.0) on a matrix in which rows represented observations and columns represented normalised features. We extracted cross-ome features (i.e., retaining only significant correlations at p<0.01) that spanned across different assays or sample types. The threshold of p<0.01 was selected to favour discovery.

#### Adjustment for multiple testing

In the current study we do not correct for multiple testing to favour discovery of novel associations for subsequent validation.

### Data visualisation

Data were visualised in R. Boxplots, scatter plots, and bubble plots, or where otherwise specified below, were visualised using ggplot2 (version 3.5.0). Principal component analysis was conducted using the FactoMineR ^86^ T package (version 2.9) and coordinates extracted for screeplots, while biplots were visualised using factoextra ^87^ (version 1.0.7). Heatmaps were generated using ComplexHeatmap ^88^ (version 2.12.1). Tree diagrams for immune populations were generated by transforming hierarchical data into a tree format using the tidytree package (version 0.4.6) and visualising the tree using the ggtree package (version 3.10.1). The Chord diagram for cross-ome correlations were visualised using the circlize R package (version 0.4.16). Cross-ome network diagrams were generated using the igraph R package (version 2.0.2), filtering first connections of selected biomarkers. Long format data were transformed into a network using the graph_from_edgelist function. Node colours were set by assay type while edge colour and size was plotted by correlation sign and strength.

#### Acknowledgements

We thank all participants of the TirolGESUND study and individuals involved in the administration of the study. We thank Nora Gibitz-Eisath for assistance with conducting urine cotinine quantitation. We thank our research nurses Gabi Hilber, Barbara Rovara and Birgit Kröss. We thank Maria Meister and her lab team for the assistance with sample processing. We thank the the vascular measurement team Silvia Komarek, Benjamin Dejakum, Alex Messner (all Department of Neurology, Medical University of Innsbruck; VASCage - Research Centre on Vascular Ageing and Stroke, Innsbruck), Bernhard Winder(Department of Vascular Surgery, Feldkirch Hospital, Feldkirch, Austria), and Johannes Nairz (VASCage- Research Centre on Vascular Ageing and Stroke, Innsbruck; Department of Pediatrics II and III, Medical University of Innsbruck). We thank Ute Pichler, Markus Augschöll, David Ebner who performed dietetic counselling for smoking cessation participants. We thank the psychology students (Minou Mohraz, Olaya Roces Sanchez, Rosa Huber, Carina Zeitler, Anne Harbring, Rosa Ottmann, Caroline Siebert, Valeria Kovalchuk, Elvira Galeazzo, Lina Förster, Pauline Raßbach, Sonja Breu, Anna Zellmer, Sonja Breu, Elif Algül, Melisa Amin, Johanna Haessler, Lilli Schulz, Valerie Nickel, Lea Rosenstock) for coaching and supporting the volunteers.

## Author contributions [CRediT statement]

**Chiara Maria Stella Herzog**: conceptualisation, methodology, formal analysis, data curation, writing - original draft, visualisation, supervision, project administration.

**Charlotte Dafni Vavourakis**: methodology, formal analysis, visualisation, data curation, writing - original draft.

**Bente Theeuwes**: formal analysis, visualisation, data curation, software, writing - review and editing.

**Elisa Redl**: investigation, project administration, writing - review and editing.

**Christina Watschinger**: investigation, project administration, writing - review and editing.

**Gabriel Knoll**: investigation, data curation, writing - review and editing

**Magdalena Hagen**: investigation, data curation, writing - review and editing

**Andreas Haider**: investigation, validation, writing - review and editing

**Hans-Peter Platzer**: investigation, data curation, writing - review and editing

**Umesh Kumar**: investigation, data curation, writing - review and editing

**Sophia Zollner-Kiechl**: investigation, data curation, writing - review and editing

**Michael Knoflach**: investigation, data curation, writing - review and editing

**Nora Gibitz-Eisath**: investigation, data curation

**Stefan Öhler**: investigation, supervision

**Verena Lindner**: investigation, supervision

**Anna Wimmer**: investigation, supervision Tobias Greitemeyer: investigation, supervision

**Peter Widschwendter**: investigation, supervision

**Sonja Sturm**: investigation, supervision, data curation, writing - review and editing

**Hermann Stuppner**: resources, writing - review and editing

**Birgit Weinberger**: supervision, resources, writing - review and editing

**Alexander Moschen**: resources, project administration, writing - review and editing

**Alex Höller**: supervision, resources

**Wolfgang Schobersberger**: supervision, resources

**Christian Haring:** supervision, resources

**Martin Widschwendter**: conceptualisation, project administration, resources, writing - original draft, supervision, funding acquisition.

## Data availability

Data availability is described in detail in a corresponding data description paper ^12^ and links to digital object identifiers of datasets will be provided on the Tyrol Lifestyle Atlas Data Portal (https://eutops.github.io/lifestyle-atlas/). In brief, methylation and microbiome data are deposited on EGA under accession codes EGAS00001007841 and EGAS00001007843 [to be released with publication]. Other assay data are deposited on Zenodo in tabular format. Anonymised data will be available openly after a one year embargo period. Study data can be explored using the Tyrol Lifestyle Atlas Data App (https://eutops.github.io/lifestyle-atlas/docs/explore/app.html).

## Code availability

Code for analyses is provided on github under (https://github.com/chiaraherzog/MultiOmics_SmokingCessation).

