## Supporting information for "Longitudinal multi-omic evaluation of biomarkers of health and ageing over smoking cessation intervention"

**Extended Data Tables** **3**
Extended Data Table 1. Baseline characteristics of participants included in the study. 3
Extended Data Table 2. Intention to treat analysis comparing baseline values to month 2, 4, and 6 (paired two-sided Wilcoxon test) in individuals who completed the study. 3
Extended Data Table 3. Intention to treat analysis using linear mixed effects models to compare baseline values to month 2, 4, and 6, adjusting for age at baseline and smoking pack years in individuals who completed the study. 3
Extended Data Table 4. Comparison of changes from baseline in the higher compliance versus lower compliance group (unpaired two-sided Wilcoxon test at month 2, 4, and 6) in individuals who completed the study. 3
Extended Data Table 5. Linear mixed effects models exploring the interaction of time and high compliance in individuals who completed the study, adjusting for age at consent and smoking pack years. 3
Extended Data Table 6. Per protocol analysis comparing baseline values to month 2, 4, and 6 (paired two-sided Wilcoxon test) in highly compliant individuals who completed the study. 3
Extended Data Table 7. Per protocol analysis assessing the impact of time in highly compliant individuals who completed the study using linear mixed-effects model, adjusting for age at consent and smoking pack years. 4
Extended Data Table 8. Overall and pairwise permutational analysis of variance of the Aitchison distance for groups with different smoking status at baseline and months 2, 4 and 6 of the smoking cessation intervention. 5
Extended Data Table 9. Associations of baseline features with smoking cessation success and buccal methylation change at month 6. 5
Extended Data Table 10. Repeated measures correlation of key variables. 5
**Extended Data Figures** **6**
Extended Data Figure 1. Association of principal components 1 and 2 of top variable features with age and smoking behaviour. 6
Extended Data Figure 2. Association of immune cell populations with smoking at baseline and detailed population changes over time. 7
Extended Data Figure 3. Principal components of epigenetic data and association with cell composition, validation of smoking signatures, modules of DNAmGrimAgeV2 over time, and exploration of changes with nicotine replacement tools. 9
Extended Data Figure 4. Principal components of metabolites in saliva and urine and their association with cigarettes at baseline. 11
Extended Data Figure 5. Microbiome analysis details. 12
Extended Data Figure 6. Integrative omic MEFISTO analyses. 13
Extended Data Figure 7. Integration analysis examples of multi-omic repeated measures correlations. 14
**Supplementary Data** **15**
Supplementary Data Figure 1. Evaluation of the ImmuneSMK predictor. 15

### Extended Data Tables

**Extended Data Table 1. Baseline characteristics of participants included in the study.**

| Characteristic | Overall<br>n = 42 | Cessation (M6)<br>n = 14 | No cessation (M6)<br>n = 10 | Dropout<br>n = 18 |
|---|---|---|---|---|
| **Age at consent, Mean (SD)** | 43.2 (6.9) | 43.3 (6.3) | 44.3 (8.3) | 42.5 (6.8) |
| **BMI at consent ($kg/m^2$), Mean (SD)** | 25.3 (5.4) | 24.6 (5.9) | 26.5 (5.7) | 25.2 (5.1) |
| **Postmenopausal, n (%)** | 8 (19) | 3 (21) | 3 (30) | 2 (11) |
| **Cigarettes per day at baseline, Mean (SD)** | 16.9 (5.4) | 14.9 (4.0) | 15.6 (3.5) | 19.3 (6.4) |
| **Smoking pack years, Median (IQR)** | 20.0 (10.2) | 18.0 (8.0) | 18.8 (8.0) | 23.0 (8.9) |
| **Weekly intense activity (min), n (%)** | | | | |
| no intense activity | 23 (55) | 9 (64) | 4 (40) | 10 (56) |
| 30-149 min | 10 (24) | 3 (21) | 2 (20) | 5 (28) |
| ≥150 min | 7 (17) | 2 (14) | 3 (30) | 2 (11) |
| unknown | 2 (4.8) | 0 (0) | 1 (10) | 1 (5.6) |
| **Systolic blood pressure (mmHg), Mean (SD)** | 114.8 (13.4) | 113.2 (13.5) | 109.5 (8.0) | 119.1 (15.0) |
| **Diastolic blood pressure (mmHg), Mean (SD)** | 74.3 (9.0) | 73.6 (7.2) | 71.5 (3.4) | 76.5 (12.0) |
| **$VO_2$peak (mL/kg/min), Mean (SD)** | 33.8 (8.6) | 34.5 (7.6) | 35.6 (10.9) | 32.1 (8.0) |
| **Total cholesterol (mg/dL), Mean (SD)** | 187.7 (32.0) | 185.5 (30.6) | 204.6 (34.2) | 179.6 (29.9) |
| **Triglycerides (mg/dL), Median (IQR)** | 99.0 (85.0) | 99.5 (67.5) | 72.0 (64.3) | 112.0 (99.0) |
| **HbA1c (%), Mean (SD)** | 5.3 (0.2) | 5.3 (0.3) | 5.3 (0.1) | 5.3 (0.3) |
| **Fasting glucose (mg/dL), Mean (SD)** | 83.2 (17.6) | 79.6 (10.2) | 74.3 (19.5) | 91.4 (18.6) |
| **Haemoglobin (g/dL), Mean (SD)** | 13.4 (0.9) | 13.3 (1.0) | 13.2 (0.7) | 13.6 (0.9) |
| **Erythocytes per mL, Mean (SD)** | 4.4 (0.4) | 4.4 (0.3) | 4.4 (0.3) | 4.5 (0.4) |

**Extended Data Table 2. Intention to treat analysis comparing baseline values to month 2, 4, and 6 (paired two-sided Wilcoxon test) in individuals who completed the study.**
Provided as .xlsx file.

**Extended Data Table 3. Intention to treat analysis using linear mixed effects models to compare baseline values to month 2, 4, and 6, adjusting for age at baseline and smoking pack years in individuals who completed the study.**
Each variable was normalised to provide standardised estimate sizes. Time was coded as a categorical variable (visitId). Provided as .xlsx file.

**Extended Data Table 4. Comparison of changes from baseline in the higher compliance versus lower compliance group (unpaired two-sided Wilcoxon test at month 2, 4, and 6) in individuals who completed the study.**
Provided as .xlsx file.

**Extended Data Table 5. Linear mixed effects models exploring the interaction of time and high compliance in individuals who completed the study, adjusting for age at consent and smoking pack years.**
Each variable was normalised to provide standardised estimate sizes. Time was coded as a categorical variable (visitId). Provided as .xlsx file.

**Extended Data Table 6. Per protocol analysis comparing baseline values to month 2, 4, and 6 (paired two-sided Wilcoxon test) in highly compliant individuals who completed the study.**
Provided as .xlsx file.

**Extended Data Table 7. Per protocol analysis assessing the impact of time in highly compliant individuals who completed the study using linear mixed-effects model, adjusting for age at consent and smoking pack years.**

Each variable was normalised to provide standardised estimate sizes. Time was coded as a categorical variable (visitId). Provided as .xlsx file.

**Extended Data Table 8. Overall and pairwise permutational analysis of variance of the Aitchison distance for groups with different smoking status at baseline and months 2, 4 and 6 of the smoking cessation intervention.**

| Sample type | Comparison | P value | Adjusted p value |
|---|---|---|---|
| **Saliva** | Smoker baseline vs Never smoker (control) | 0.001 | NA |
| **Saliva** | M2 no smoking cessation vs M2 smoking cessation vs Never smoker (control) | 0.0285 | NA |
| **Saliva** | Never smoker (control) vs M2 no smoking cessation | 0.002 | 0.006 |
| **Saliva** | Never smoker (control) vs M2 smoking cessation | 0.545 | 1 |
| **Saliva** | M2 no smoking cessation vs M2 smoking cessation | 0.503 | 1 |
| **Saliva** | M4 no smoking cessation vs M4 smoking cessation vs Never smoker (control) | 0.0246 | NA |
| **Saliva** | Never smoker (control) vs M4 no smoking cessation | 0.009 | 0.027 |
| **Saliva** | Never smoker (control) vs M4 smoking cessation | 0.094 | 0.282 |
| **Saliva** | M4 no smoking cessation vs M4 smoking cessation | 0.644 | 1 |
| **Saliva** | M6 no smoking cessation vs M6 smoking cessation vs Never smoker (control) | 0.0024 | NA |
| **Saliva** | Never smoker (control) vs M6 no smoking cessation | 0.001 | 0.003 |
| **Saliva** | Never smoker (control) vs M6 smoking cessation | 0.241 | 0.723 |
| **Saliva** | M6 no smoking cessation vs M6 smoking cessation | 0.028 | 0.084 |
| **Stool** | Smoker baseline vs Never smoker (control) | 0.0528 | NA |
| **Stool** | M2 no smoking cessation vs M2 smoking cessation vs Never smoker (control) | 0.2564 | NA |
| **Stool** | Never smoker (control) vs M2 no smoking cessation | 0.333 | 0.999 |
| **Stool** | Never smoker (control) vs M2 smoking cessation | 0.231 | 0.693 |
| **Stool** | M2 no smoking cessation vs M2 smoking cessation | 0.357 | 1 |
| **Stool** | M4 no smoking cessation vs M4 smoking cessation vs Never smoker (control) | 0.1629 | NA |
| **Stool** | Never smoker (control) vs M4 no smoking cessation | 0.442 | 1 |
| **Stool** | Never smoker (control) vs M4 smoking cessation | 0.078 | 0.234 |
| **Stool** | M4 no smoking cessation vs M4 smoking cessation | 0.445 | 1 |
| **Stool** | M6 no smoking cessation vs M6 smoking cessation vs Never smoker (control) | 0.116 | NA |
| **Stool** | Never smoker (control) vs M6 no smoking cessation | 0.171 | 0.513 |
| **Stool** | Never smoker (control) vs M6 smoking cessation | 0.232 | 0.696 |
| **Stool** | M6 no smoking cessation vs M6 smoking cessation | 0.106 | 0.318 |

**Extended Data Table 9. Associations of baseline features with smoking cessation success and buccal methylation change at month 6.**
Provided as .xlsx file.

**Extended Data Table 10. Repeated measures correlation of key variables.**
Provided as .xlsx file.

### Extended Data Figures

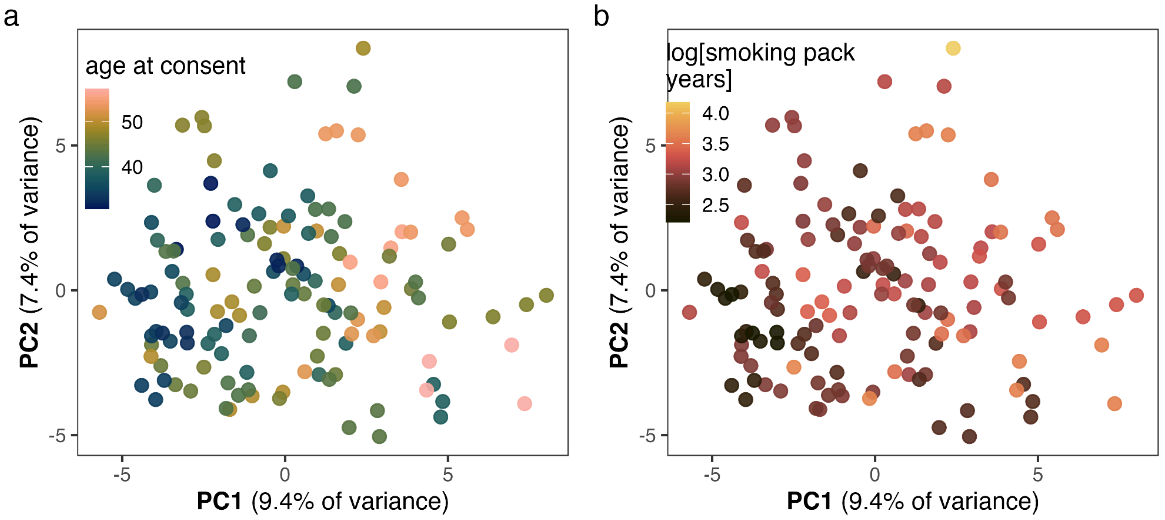


**Extended Data Figure 1. Association of principal components 1 and 2 of top variable features with age and smoking behaviour.**

**a** Principal components 1 and 2 of top 30% variable features coloured by age and **b** smoking pack years (log).

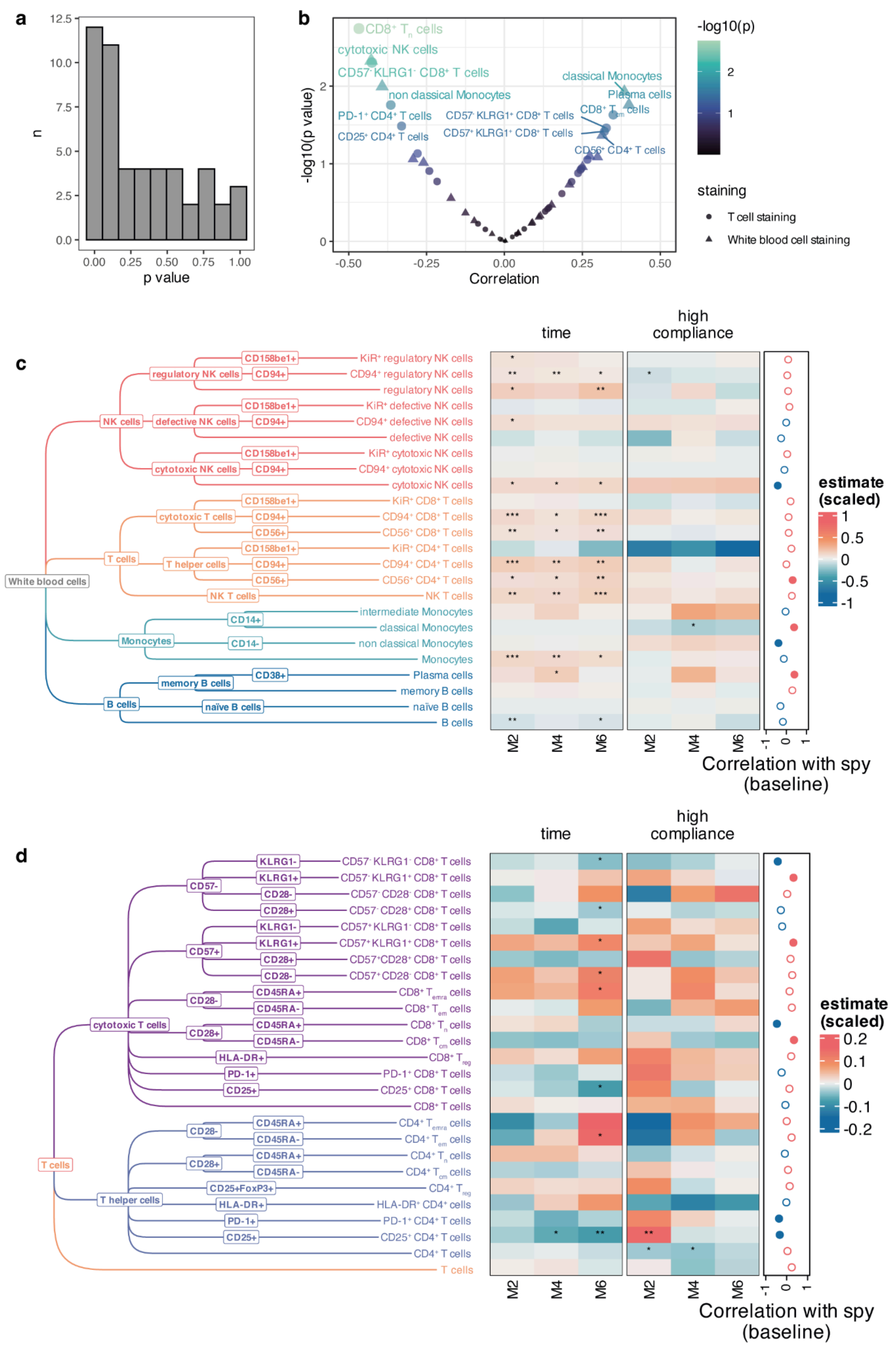


**Extended Data Figure 2. Association of immune cell populations with smoking at baseline and detailed population changes over time.**

**a** Histogram of p values for correlation of immune cell proportion abundance and smoking pack years at baseline. **b** Volcano plot of correlation of immune cell populations with smoking pack years at baseline. **c** Tree diagram for immune cell populations analysed using white blood cell staining and estimates from linear-mixed effects models impact of time (value ~ age at consent + smoking pack years at consent + visitId + (1|subjectId)), interaction between time and high compliance (value ~ age at consent + smoking pack years at consent + visitId*compliance + (1|subjectId)). *, **, *** indicate p <0.05, <0.01, and <0.001, respectively. Correlation with age at baseline smoking pack year values is indicated on the right hand side (positive or negative in red and blue, respectively). Significant correlations at p<0.05 are indicated with filled circles. **e** Tree diagram for immune cell populations analysed using T cell staining and estimates from linear-mixed effects models impact of time (value ~ age at consent + smoking pack years at consent + visitId + (1|subjectId)), interaction between time and high compliance (value ~ age at consent + smoking pack years at consent + visitId*compliance + (1|subjectId)). *, **, *** indicate p<0.05, <0.01, and <0.001, respectively. Correlation with age at baseline smoking pack year values is indicated on the right hand side (positive or negative in red and blue, respectively). Significant correlations at p<0.05 are indicated with filled circles.

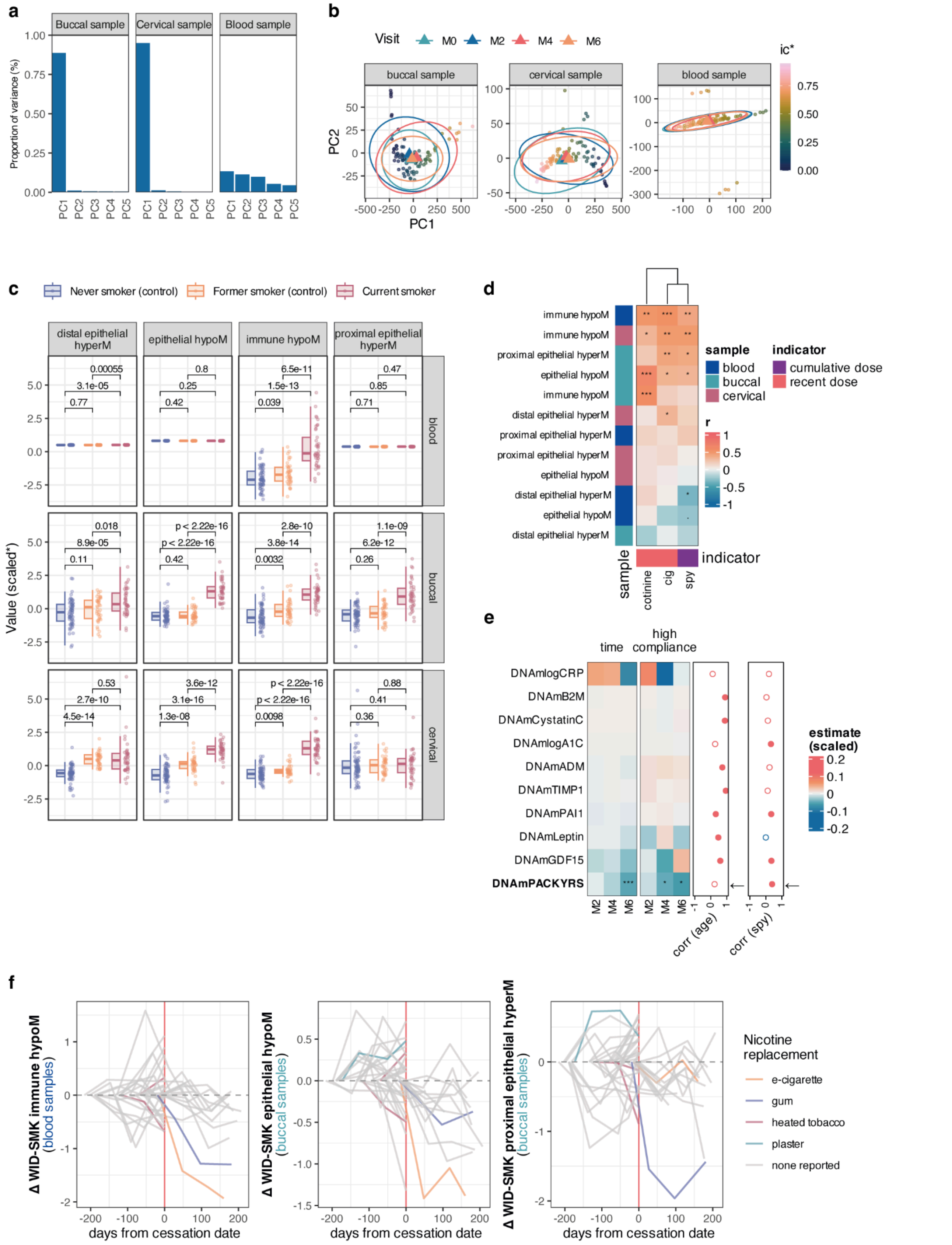

**Extended Data Figure 3. Principal components of epigenetic data and association with cell composition, validation of smoking signatures, modules of DNAmGrimAgeV2 over time, and exploration of changes with nicotine replacement tools.**

**a** Scree plot of first 5 principal components in buccal, cervical, and blood data. **b** Scatterplot of principal components 1 and 2 in buccal, cervical, and blood samples, coloured by immune cell composition (neutrophil proportion in blood). Large signs indicate centroids of samples for each visit. **c** Validation of previously-described cell-specific smoking signatures using baseline samples of the current study and a parallel study that contains never and former smokers with several quit-years. **d** Association of smoking signatures in each tissue at baseline with cotinine value, cigarettes per day at baseline (cig) or smoking pack years (spy). **e** LME estimates of GrimAgeV2 modules over time. **f** Changes with nicotine replacement and timing of smoking. Samples are plotted relative to the time from cessation date in days, with samples collected at visits prior to cessation shown as negative. Individuals who did not give up smoking are plotted as having all visits pre smoking cessation, where day 0 is the final visit. Change values are shown. For the individual with nicotine use, no month 0 blood sample was available and hence no change is shown for them for WID-SMK immune hypoM.

*, p<0.05, **, p<0.01, ***, p<0.001, ****, p<0.0001, in unpaired *t* Tests (**c**), Pearson's correlation test (**d**), or LME (**e**).

**Abbreviations**: cig, cigarettes per day at baseline. spy, smoking pack years at baseline.

Box plots correspond to standard Tukey representation, with boxes indicating median and interquartile ranges and whiskers indicating ± 1.5 times interquartile ranges. Individual data points are shown where possible.

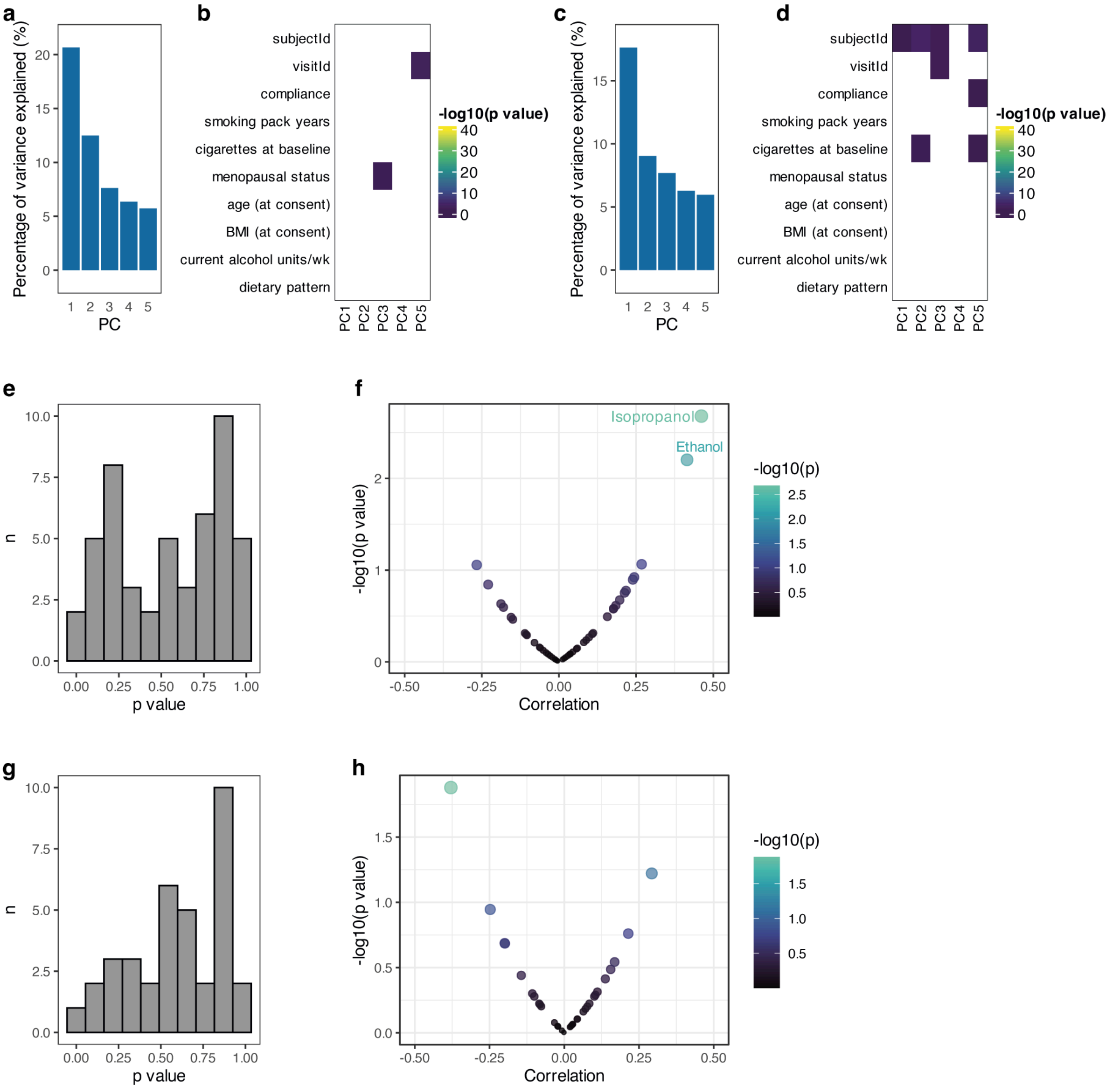


**Extended Data Figure 4. Principal components of metabolites in saliva and urine and their association with cigarettes at baseline.**

**a** Scree diagram of the first five principal components of saliva metabolome data. **b** Heatmap of significant associations of covariates with the first five principal components of saliva metabolome data. **c** Scree diagram of the first five principal components of urine metabolome data. **b** Heatmap of significant associations of covariates with the first five principal components of urine metabolome data. **e** Histogram of p values for the association of reported cigarettes per day at baseline and baseline saliva metabolome values. **f** Volcano plot of spearman correlation value and -log10(p value) for the correlation of reported cigarettes per day at baseline and baseline saliva metabolome values. Associations with $p<0.05$ are labelled. **g** Histogram of p values for the association of reported cigarettes per day at baseline and baseline urine metabolome values. **h** Volcano plot of spearman correlation value and -log10(p value) for the correlation of reported cigarettes per day at baseline and baseline urine metabolome values. No associations were significant at $p<0.05$.

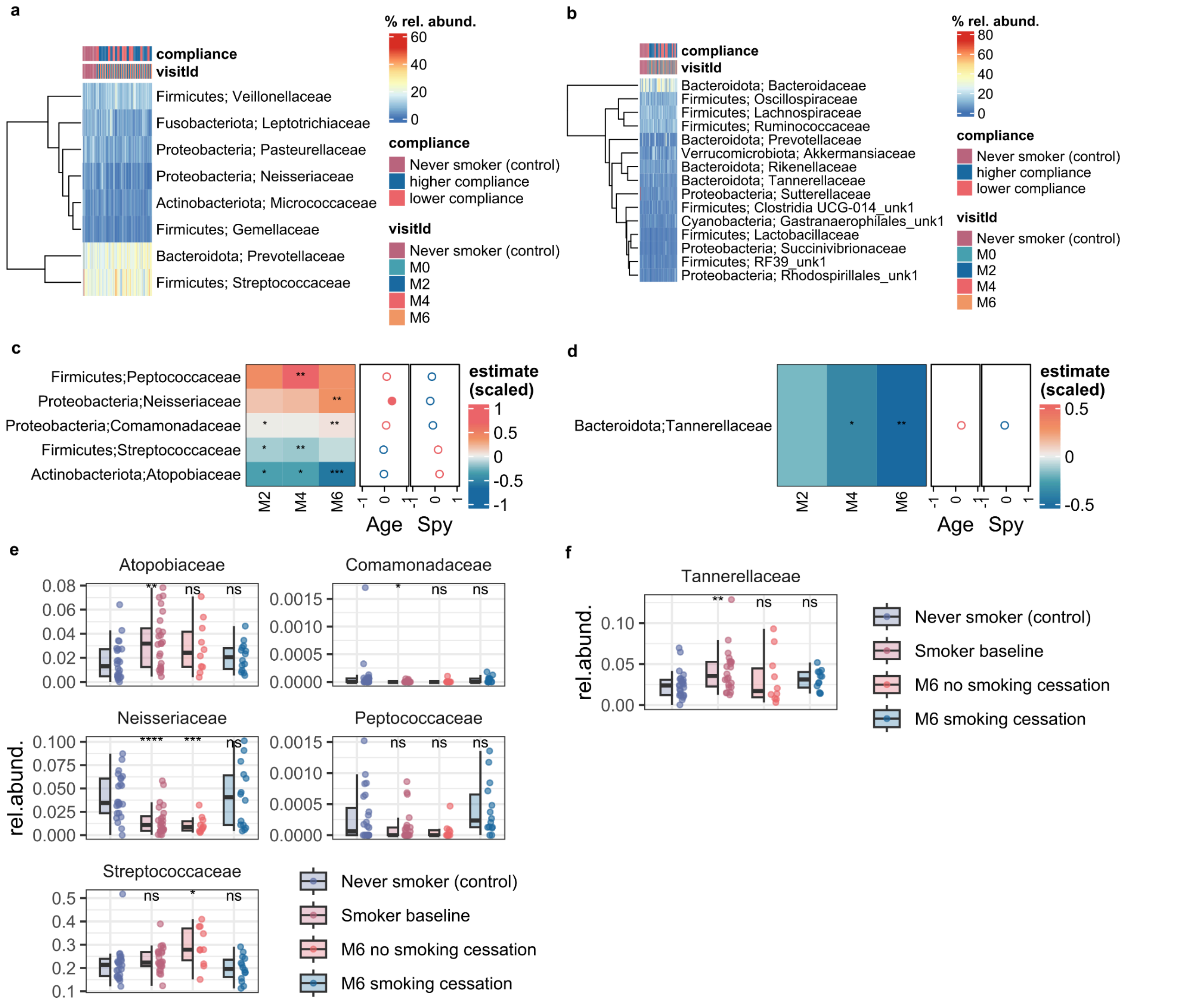


**Extended Data Figure 5. Microbiome analysis details.**

**a** Relative abundance of the most abundant microbial families (minimum 10% relative abundance in any sample) in the saliva and **b** stool data sets, respectively. **c** Time estimates for the microbial families that significantly changed (p value ≤ 0.01) at month 2, 4 or 6 compared with baseline in the higher compliance smoking cessation group in the oral and **d** gut microbiomes, respectively. Linear mixed models were corrected for age and smoking pack years at consent, for which Pearson correlation with relative abundance at baseline for the respective families is depicted as well (full circles p values p value < 0.05). **e** Relative abundance of the differentially abundant microbial families upon successful smoking cessation in never-smokers, smokers at baseline and month 6, and former-smokers at month 6 in saliva and **f** stool samples.

Significance levels in c-f are denoted as follows: ns = non significant; * = p value<0.05; ** p value<0.01, *** = p value<0.001).

Box plots correspond to standard Tukey representation, with boxes indicating median and interquartile ranges and whiskers indicating ± 1.5 times interquartile ranges. Individual data points are shown where possible.

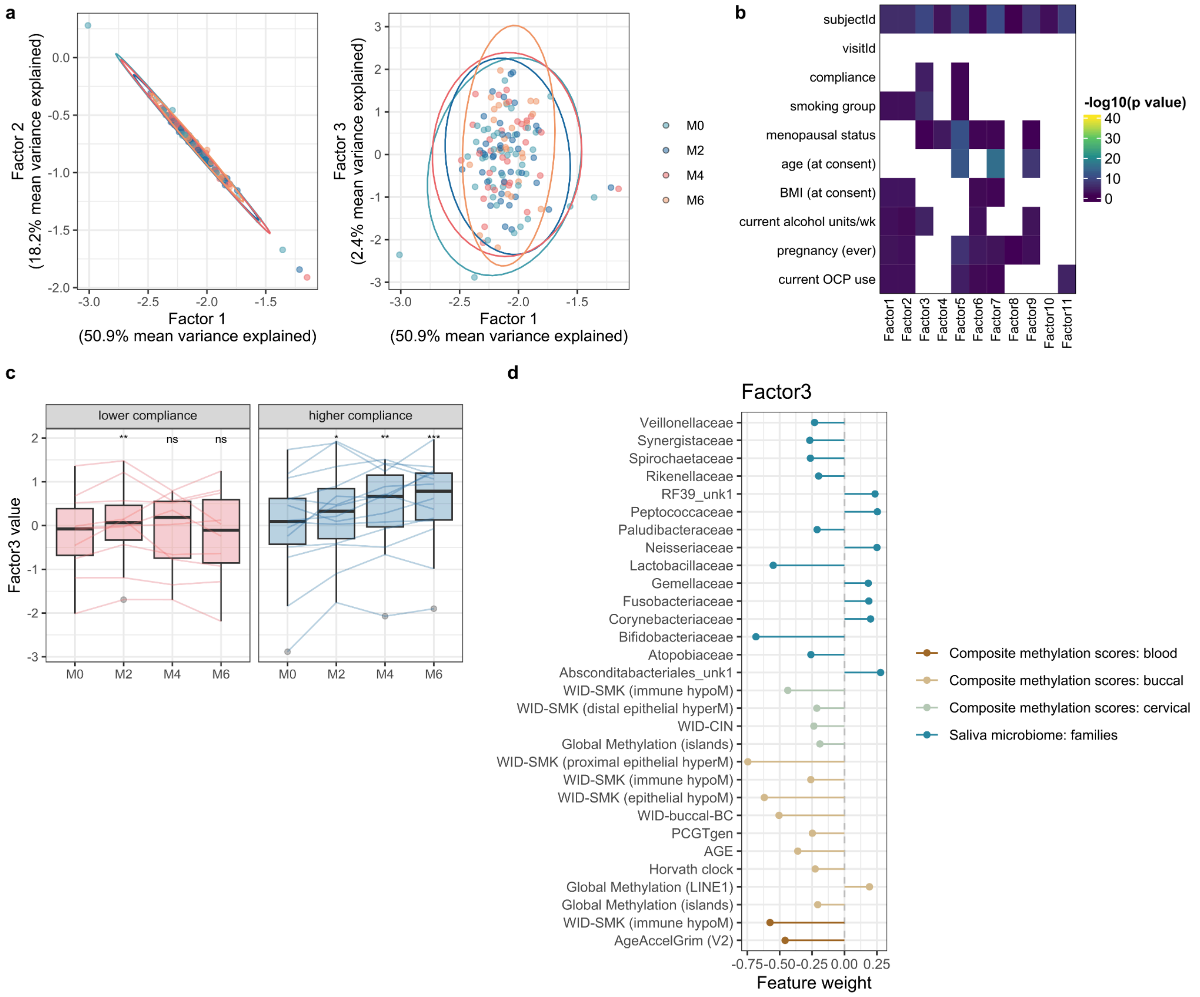


**Extended Data Figure 6. Integrative omic MEFISTO analyses.**

**a** Biplots depict the factor values for the top 3 latent factors found by MEFISTO across the epigenomes (composite methylation score and immune age), metabolomes, microbiomes, immune cell cell stainings, and blood haemogram. The mean of the variances explained in each omic dataset by the respective factors is given on the x, and y axes, and samples are coloured by visitId. **b** Heatmap showing the significance of the 11 detected latent factors with recorded covariates. **c** The grouped bar plot depicts the changes over time of latent factor 3 for the two compliance groups. Significance changes over time for each subgroup was verified with paired Wilcoxon tests (ns, *, **, ***, **** denote p≥0.05, <0.05, <0.01, <0.001, and <0.0001, respectively). **d** Weights of the top 30 features associated with latent factor 3.

Box plots correspond to standard Tukey representation, with boxes indicating median and interquartile ranges and whiskers indicating ± 1.5 times interquartile ranges. Individual data points are shown where possible.

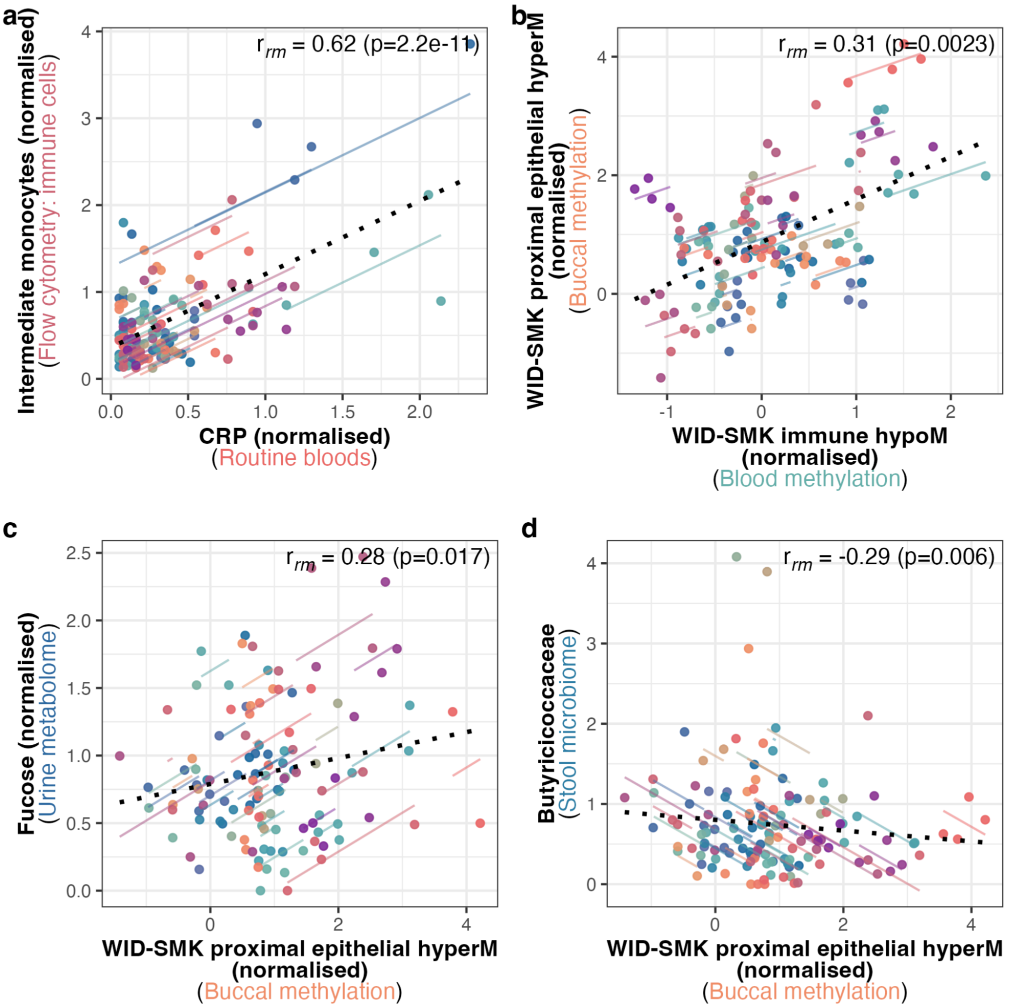


**Extended Data Figure 7. Integration analysis examples of multi-omic repeated measures correlations.**

**a** Correlation of CRP with intermediate monocyte counts. **b** Repeated measures correlation of buccal WID-SMK proximal hyperM values with blood WID-SMK immune hypoM values, **c** Urine fucose, or **d** stool Butyricicoccaceae. Points are coloured by individuals, with model fits for each individual shown. Overall correlation line is shown in a dotted line. Correlation R and p values are obtained from repeated measures correlation.

### Supplementary Data

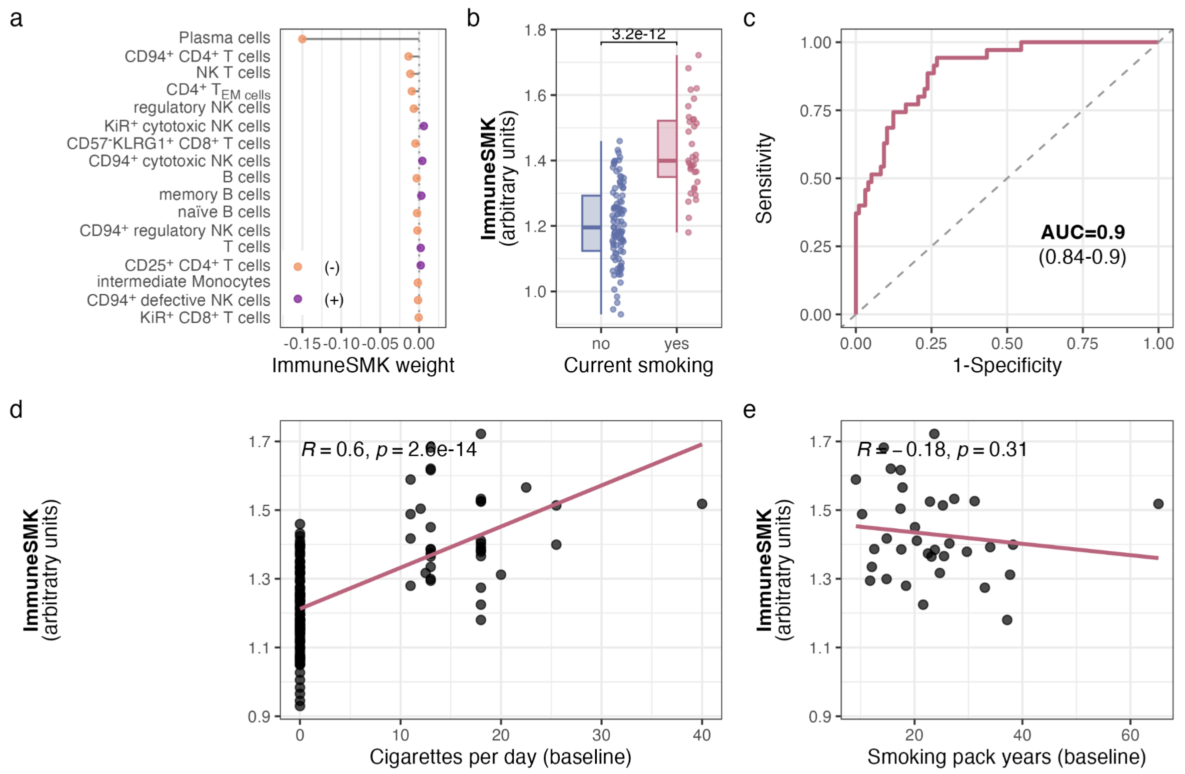


**Supplementary Data Figure 1. Evaluation of the ImmuneSMK predictor.**

**a** Weights of the ImmuneSMK predictor. **b** ImmuneSMK values by current smoking at baseline. p value is derived from an unpaired two-sided Wilcoxon tests. **c** Area under the curve (AUC) of the ImmuneSMK predictor. **d** ImmuneSMK predictor by cigarettes per day at baseline. Correlation and p value are derived from Spearman correlation test. **e** ImmuneSMK predictor by smoking pack years in smokers at baseline. Correlation and p value are derived from Spearman correlation test.
